# Ribosome Quality Control Mitigates Proteotoxic Stress in Aneuploid Cells

**DOI:** 10.64898/2026.01.19.700285

**Authors:** Sonia Viganò, Marica R. Ippolito, Simone Scorzoni, Lara Barrio, Alessia Loffreda, Marina Rodriguez-Muñoz, Deborah Donzel, Yonatan Eliezer, Laura Pontano Vaites, Joao A. Paulo, Mattia Marenda, Alessandro Palma, Uri Ben-David, Gabriella Viero, Marco Milán, Stefano Santaguida

## Abstract

Aneuploidy is widespread in tumors, but how cancer cells adapt to aneuploidy-induced cellular stresses remains poorly understood. Here, we focus on the mechanisms employed to cope with proteostasis disruption, a major stress caused by aneuploidy. We show that aneuploid cells exhibit a significant accumulation of ribosomes enclosed within autophagosomes, ultimately degraded through lysosome-mediated processes. Our data also indicate that limited folding capacity of newly synthesized polypeptides leads to lysosome-mediated degradation of ribosomes. We also found that the E3 ligase ZNF598 marks these ribosomes for degradation, thus clearing translationally-impaired ribosomes. Importantly, highly aneuploid tumors display a positive correlation with ZNF598 expression, while being negatively associated with ribosomal signatures. This suggests that ribosome-associated quality control is crucial for cancer cell survival under proteotoxic stress. Our study uncovers molecular events in response to proteotoxic stress in aneuploid cells and suggests that components of ribosome-associated quality control, including ZNF598, could serve as promising targets in cancer therapy.

## INTRODUCTION

Aneuploidy - a condition in which a cell harbours an incorrect chromosome number - is a widespread feature of tumours(*1–4*). Aneuploidy-induced genome and proteome imbalances have several consequences on cellular homeostasis and lead to impaired proliferation(*5–8*), increased DNA damage and genome instability(*9–13*) - which fuel the evolution of complex karyotypes(*14–19*) -, metabolic(*7*, *20*, *21*) and proteotoxic stresses(*3*, *4*, *22*, *23*). The latter describes the negative impact of aneuploidy on cell proteostasis, which is physiologically maintained in healthy cells through a fine balance between protein synthesis, protein folding and protein degradation(*24*). Indeed, any deviation in terms of chromosome number from the euploid karyotype has a profound impact on the stoichiometry of protein complex subunits(*22*) and overwhelms the chaperone-mediated folding machinery(*25*). These impairments are responsible for the accumulation of misfolded and unfolded proteins in aneuploid cells, as shown in yeast(*26*, *27*). In agreement with this, disrupted proteostasis has been also described in *Drosophila melanogaster* models of aneuploidy(*21*, *28*), as well as in human cells following aneuploidy induction(*4*, *29*) and in CH21 trisomic models(*30*). Along this line, aneuploid cells accumulate autophagosomes(*8*, *31*), indicating that there is an increase in the cellular cargoes engulfed in autophagic structures.

Disruption of protein homeostasis represents a major threat for cell survival(*32–34*) and multiple protein quality control pathways are impaired in aneuploid cells(*8*, *25*, *26*). Therefore, unravelling the molecular mechanisms that orchestrate the response to proteotoxic stress remain of foremost importance. A comprehensive understanding of these processes could potentially reveal vulnerabilities that can be exploited to selectively eradicate aneuploid cancer cells.

Previous studies have examined the autophagic processes in aneuploid cells and have reported an increase in autophagic cargo(*8*, *31*). However, it remains unclear which structures are encapsulated within autophagosomes, the reason why these cargoes are encapsulated, and why they are difficult to degrade.

Here, by isolating and analysing the content of autophagosomes in aneuploid cells, we found that ribosomal proteins are engulfed within them. Our data indicate that an impaired protein folding is strictly connected to autophagic recognition of translationally impaired ribosomes, immediately following chromosome mis-segregation. Further, we show that these ribosomes undergo a lysosome-mediated degradation (*i.e.*, ribophagy) through a process involving the E3-ligase ZNF598.

Aneuploidy-driven genomic instability is known to fuel a vicious cycle leading to the generation of complex karyotypes(*12*, *13*, *15*). The accumulation of such chromosomal aberrations further exacerbates proteotoxic stress associated with aneuploidy and leads to the buildup of aggregates and dysfunctional organelles. Our analysis shows that, under these conditions, aneuploid cells experience a widespread increase in autophagic cargoes, with ribosomes being among them. As a result, ribophagy becomes a bystander, making ZNF598 dispensable. Interestingly, when we stratified human tumors based on their aneuploidy scores, we found that highly aneuploid tumors show lower ribosomal signatures and higher ZNF598 expression compared to tumors with low degrees of aneuploidy. We speculate that this reflects a global attenuation of aneuploidy-induced stresses at the ribosome level, and the increased expression of ZNF598 may allow cancer cells to regulate translation of specific transcripts necessary for proliferation.

Overall, our study reveals a previously uncharacterized cascade of events triggered in aneuploid cells in response to proteotoxic stress. These findings may have important implications for cancer treatment, suggesting that targeting ribosome quality control pathways could provide a strategic approach to selectively inhibit tumor cell proliferation.

## RESULTS

### Ribosomes are degraded in aneuploid cells

Aneuploid cells accumulate autophagic cargo(*8*, *31*). However, the nature of the macromolecular structures engulfed in autophagosomes remains elusive. To shed light on the identity of these autophagic cargo, we took advantage of an established mass spectrometry-based method to identify the content of purified autophagosomes(*35*) (**Figure 1A**). To this aim, we generated aneuploid cells via induction of chromosome mis-segregation in hTERT RPE-1 cells (thereafter, RPE1), a pseudo-diploid non-transformed human cell line. We did so by inhibiting the mitotic kinase Mps1 with the inhibitor reversine (Mps1i)(*36*) and by collecting cells after 72 hours (about three cell cycles). Next, we performed proteomic analysis using stable isotope labelling by amino acids in cell culture (SILAC) coupled to a density gradient separation protocol for organelle isolation(*35*) and compared the proteome of the autophagosomes isolated from aneuploid cells with that of control sample (cells treated with vehicle DMSO) (**Figure 1A**). Gene Ontology (GO) enrichment analysis revealed the signatures of organelles previously reported as autophagy cargo (**Figure 1B**), including ER(*37*) and ribosomes(*38*). We then validated the mass spectrometry results and we did so in two different cell lines, RPE1 and HCT116 (a pseudo-diploid human colorectal carcinoma cell line). For this, we employed the pH-sensitive Keima reporter fused with proteins from different cellular compartment (**Figure 1C**). These included ribosome (RPS3-Keima for the small subunit of the ribosome and RPL28-Keima for the large subunit of the ribosome(*39*)), ER (by using RAMP4-Keima(*40*)), mitochondria (by employing COX8-Keima(*40*)) and bulk autophagy (by monitoring LDHB-Keima(*40*)). Keima is resistant to acidic lysosomal pH whereas the fused protein will be degraded. This will result in the appearance of a “processed-Keima’’ band at the molecular weight of Keima(*40*) (**Figure 1C**).

**Figure 1.**
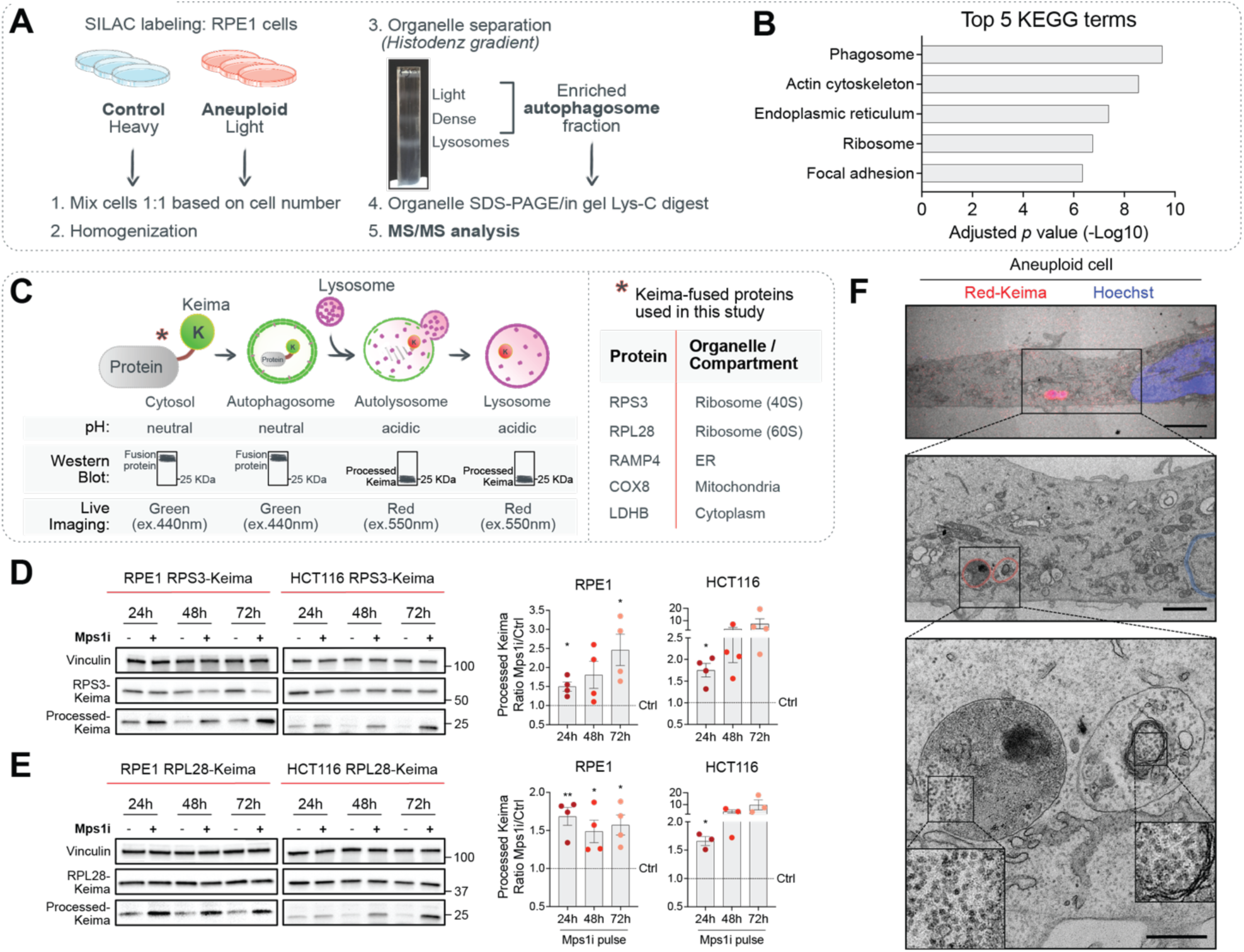
Aneuploidy leads to autophagic removal of ribosomes. **A.** Schematic illustrating the isolation of autophagosomes using a SILAC approach and subsequent mass spectrometry analysis in RPE1 cells treated with Mps1i pulse (Aneuploid) or DMSO (Control) and collected at 72 hours. **B.** Bar plot showing top 5 KEGG terms among the most enriched in autophagosomes of aneuploid cells, compared to control cells. Analysis performed using the Database for Annotation, Visualization and Integrated Discovery (DAVID) bioinformatic resource, p<0,05. **C.** Schematic indicating main features of Keima reporter and list of Keima-tagged proteins used in this study. **D.** Representative immunoblots and replicate quantitation of the indicated RPS3-Keima cell lines treated with Mps1i pulse or DMSO (control) and collected at 24, 48 or 72 hours; increased level of processed-Keima indicates ribosome degradation; vinculin was used as loading control. Mean ± SEM, *n*=4; one sample and Wilcoxon test (Mps1i at each time-point *vs* respective control=1): * indicates *p*=0.0258 (RPE1-24h) or *p*=0.0379 (RPE1-72h) or *p*=0.0170 (HCT116-24h). **E.** Representative immunoblots and replicate quantitation of the indicated RPL28-Keima cell lines treated with Mps1i pulse or DMSO (control) and collected at 24, 48 or 72 hours; increased level of processed-Keima indicates ribosome degradation; vinculin was used as loading control. Mean ± SEM, *n*=4 for RPE1, *n*=3 for HCT116; one sample and Wilcoxon test (Mps1i at each time-point *vs* respective control=1): * indicates *p*=0.0456 (RPE1-48h) or *p*=0.0230 (RPE1-72h) or *p*=0.0142 (HCT116-24h), ** indicates *p*=0.0099 (RPE1-24h). **F.** Representative images of RPE1 Ribo-Keima cell treated with Mps1i pulse and selected for CLEM: alignment and merge between confocal image and electron microscopy image (top) and magnified electron microscopy images showing ribosomes (bottom) within Red-Keima positive single-membraned autolysosomes (red profiles, centre and bottom); Hoechst was used to stain DNA (blue profiles); scale bar 5μm (top), 2μm (centre) and 500nm (bottom).

Using those reporters, we found that, in both RPE1 and HCT116 cell lines and for both ribosomal subunits, autophagic degradation of ribosomal proteins significantly increased immediately after chromosome mis-segregation (24 hours) (**Figure 1D, E**). Notably, this degradation began before the onset of degradation of other autophagic cargoes. Specifically, we did not observe a significant increase in mitophagy (**Figure S1A**), and ER-phagy showed only a slight increase 72 hours after chromosome mis-segregation (**Figure S1B**). Importantly, by employing LDHB-Keima reporter, we confirmed an increase in bulk autophagy in aneuploid cells, which correlated with the time after chromosome mis-segregation (**Figure S1C**), in agreement with previous reports(*8*, *22*). This reflects the degree of accumulation of aneuploidy-associated stresses within the cell population, increasing over time after the induction of chromosome segregation errors (**Figure S1C**). These results indicate that ribosomes are likely to be the first macromolecular structures to be engulfed in autophagosomes (**Figure S1D**). Thus, since ribosomes play a central role in protein homeostasis(*24*) and because aneuploid cells experience proteotoxic stress(*4*), we decided to investigate the autophagic removal of ribosomes upon chromosome segregation errors. First, to visualize and analyse the engulfment of ribosomes within autophagic structures with high resolution, we employed correlative light electron microscopy (CLEM) and observed that ribosomes are piled-up in single-membraned Red-Keima-positive autolysosomes (**Figure 1F** and **Figure S1G**). Second, we found that the extent of ribosome degradation increased with higher levels of aneuploidy (**Figure 2A, B**) - achieved by increasing concentrations of Mps1i(*16*) - and also increased at later time-points (**Figure 1D, E** and **Figure S1E, F**). Third, we took an orthogonal approach to generate aneuploid cells - independent of chemical inhibition of Mps1 - and confirm the autophagic degradation of ribosomes. In particular, we induced chromosome segregation errors by depleting two spindle assembly checkpoint (SAC) components, MAD2 or BUB1, and, also in this case, we found an increase in autophagic degradation of ribosomes (**Figure 2C, D**). Fourth, we took advantage of the pH-dependent excitation spectra shift of Keima(*40*) (**Figure 1C**) and measured the flux of Ribo-Keima to the acidic environment of lysosomes through live-cell imaging and observed a significant increase of Red-Keima signal in the aneuploid population (**Figure 2E**). Finally, we used an approach independent of Keima to monitor autophagy-dependent ribosome degradation, based on fluorescent tandem monomeric RFP-GFP-tagged RPS3(*40*). As GFP signal is quenched in acidic compartments, a higher RFP/GFP ratio indicates autophagosome-lysosome fusion and consequent ribosomal protein degradation. This is indeed consistent with our observations in aneuploid cells, where ribophagy was evident as early as 24 hours after chromosome mis-segregation, with a further increase after approximately three cell cycles (**Figure 2F** and **Figure S2A**). Taken together, these data indicate that ribosomes are among the first autophagic cargoes to be degraded following chromosome segregation errors, and this process correlates positively with the degree of aneuploidy.

**Figure 2.**
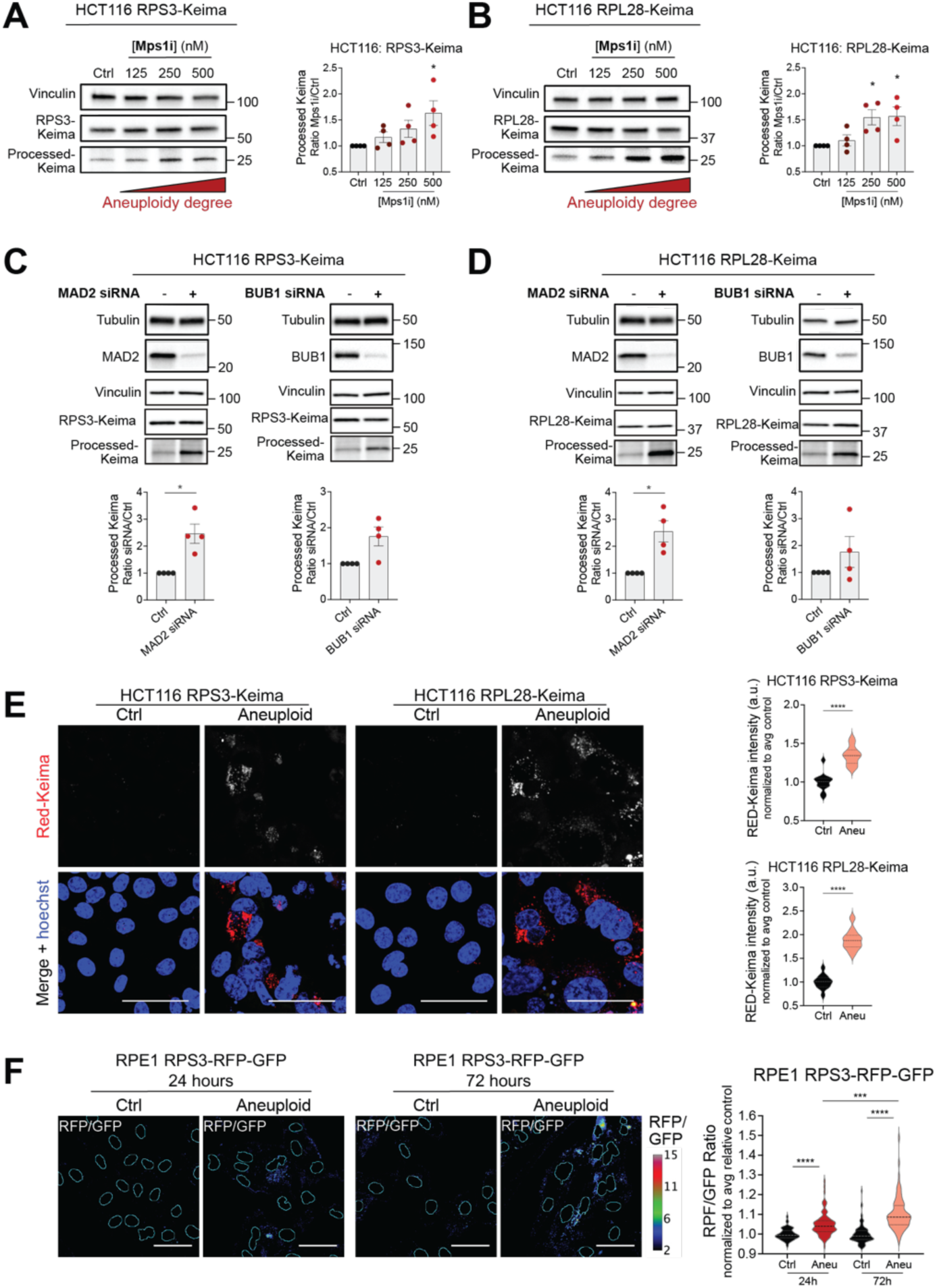
Ribosomes are degraded in aneuploid cells. **A.** Representative immunoblot and replicate quantitation of processed-Keima levels in HCT116 RPS3-Keima cells treated for 24 hours with 125nM, 250nM or 500nM Mps1i or DMSO (Ctrl); vinculin was used as loading control. Mean ± SEM, *n*=4; Kruskal-Wallis test, followed by Dunn’s multiple comparison test: * indicates *p*=0.0150. **B.** Representative immunoblot and replicate quantitation of processed-Keima levels in HCT116 RPL28-Keima cells treated for 24 hours with 125nM, 250nM or 500nM Mps1i or DMSO (Ctrl); vinculin was used as loading control. Mean ± SEM, *n*=4; Kruskal-Wallis test, followed by Dunn’s multiple comparison test: * indicates *p*=0.0329 (250nM) or *p*=0.0329 (500nM). **C.** Representative immunoblots and replicate quantitation of the HCT116 RPS3-Keima cells upon MAD2 or BUB1 siRNA (or non-targeting siRNA); increased levels of processed-Keima indicates ribosome degradation; MAD2 or BUB1 were blotted as knock-down control. Vinculin and tubulin used as loading control. Mean ± SEM, *n*=4; one sample and Wilcoxon test (each siRNA *vs* respective non-targeting siRNA=1): * indicates *p*=0.0257. **D.** Representative immunoblots and replicate quantitation of the HCT116 RPL28-Keima cells upon MAD2 or BUB1 siRNA (or non-targeting siRNA); increased levels of processed-Keima indicates ribosome degradation; MAD2 or BUB1 have been blotted as knock-down control. Vinculin and tubulin been used as loading control. Mean ± SEM, *n*=4; one sample and Wilcoxon test (each siRNA *vs* respective non-targeting siRNA=1): * indicates *p*=0.0292. **E.** Representative live-cell images and replicate quantitation of indicated HCT116 Ribo-Keima cells lines treated with Mps1i pulse or DMSO (Ctrl) and collected at 24 hours. Increased Red-Keima intensity indicates ribosome degradation; Hoechst was used to stain DNA; scale bars, 50μm. Upper quartile, lower quartile and median of each violin plot are shown, *n*=12 fields of view; unpaired Student’s t-test: **** indicates *p*<0.0001. **F.** Representative images and replicate quantitation of RFP/GFP ratio in RPE1 RPS3-RFP-GPF cells treated with Mps1i pulse or DMSO (Ctrl) and collected at 24 or 72 hours. RFP/GFP ratio was calculated as explained in Methods and visualised with a rainbow lookup table (calibration bar shown); cyan masks indicate primary nuclei; scale bars 50μm. Upper quartile, lower quartile and median of each violin plot are shown; n=3 biological replicates (30 fields of view each); Kruskal-Wallis test, followed by Dunn’s multiple comparison test: *** indicates *p*=0.0002, **** indicates *p*<0.0001.

### Ribosomes are cleared via lysosomes in aneuploid cells

Autophagy is an evolutionarily conserved catabolic pathway in which autophagic cargoes are degraded in the acidic lysosomal compartment(*41*). To assess if the removal of ribosomes in aneuploid cells relies on the canonical events of the autophagic degradation pathway, we modulated autophagy at different stages(*11*, *40*, *42*). First, we blocked the initial steps of the autophagic pathway with SAR405, an inhibitor of VPS34, which is involved in phagophore nucleation and autophagosome formation(*42*). This treatment decreased the levels of processed Keima in aneuploid cells, without affecting controls (**Figure S2B**). In addition, to interfere with the final stages of autophagy, we treated cells with Bafilomycin A_1_, an inhibitor of lysosome acidification and autophagosome-lysosome fusion(*42*). The significant decrease of processed Keima (**Figure S2C**), obtained upon Bafilomycin A_1_ treatment, confirmed the dependency of this process on lysosomal degradation in aneuploid cells. In agreement with this finding, we detected a significant reduction of Red-Keima puncta (**Figure 3A**) and RFP/GFP ratio (**Figure 3B**) in our live-cell imaging experiments, upon Bafilomycin A_1_ treatment. Overall, these results show that, in aneuploid cells, degradation of both small and large ribosomal subunits is lysosome-dependent and takes place through the canonical steps of autophagy.

**Figure 3.**
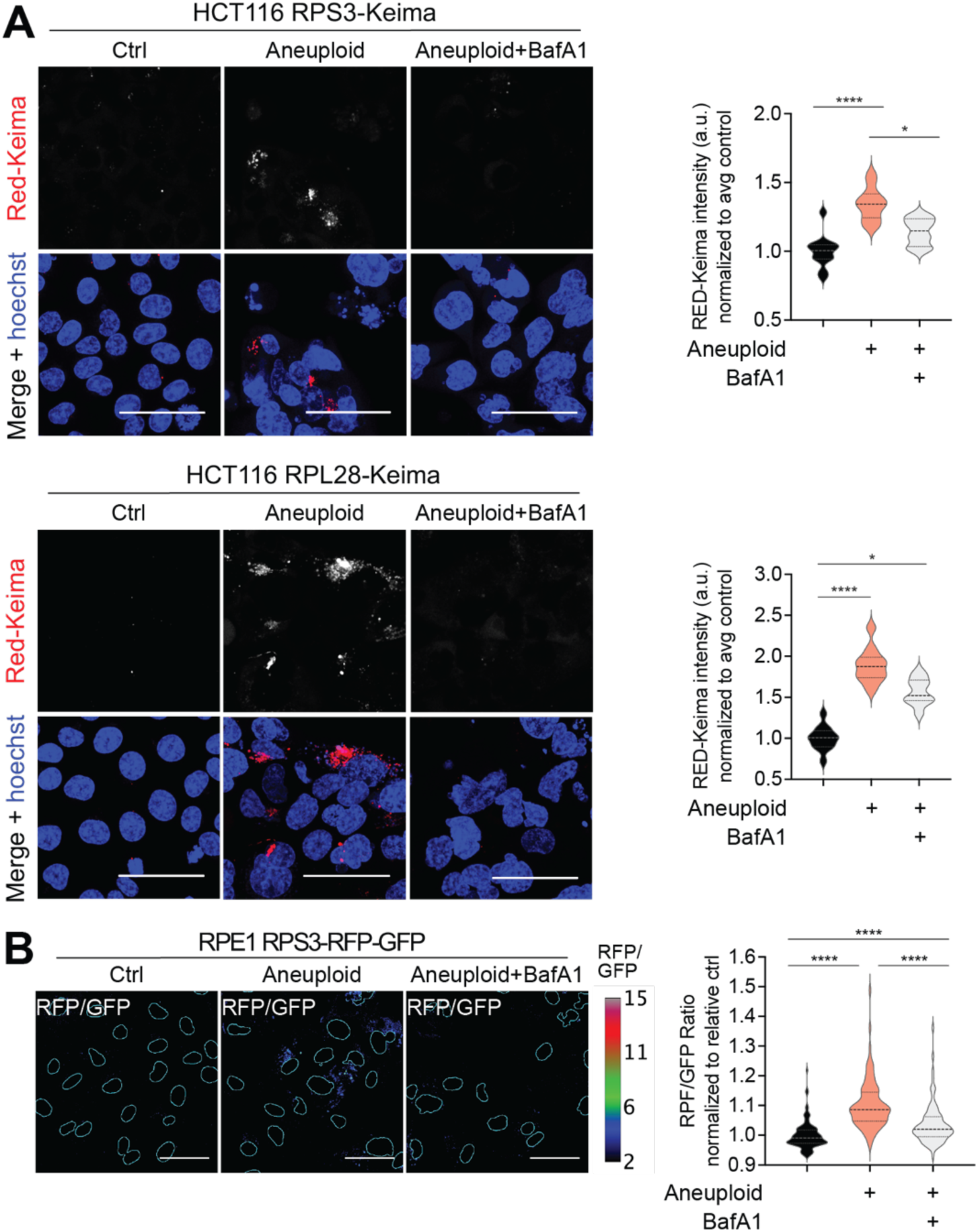
Ribosome clearance in aneuploid cells relies on canonical lysosomal degradation. **A.** Representative live-cell images and replicate quantitation of indicated HCT116 Ribo-Keima cells lines treated with Mps1i pulse or DMSO (Ctrl), collected at 24 hours and upon Bafilomycin A_1_ (BafiloA_1_) treatment (100nM, 6h). Increased Red-Keima intensity indicates ribosome degradation; Hoechst was used to stain DNA; images have been obtained from the experiment shown in Fig2E; scale bars, 50μm. Upper quartile, lower quartile and median of each violin plot are shown, *n*=12 fields of view; Kruskal-Wallis test, followed by Dunn’s multiple comparison test: * indicates *p*=0.0252 (HCT116 RPS3-Keima) or *p*=0.0150 (HCT116 RPL28-Keima), **** indicates *p*<0.0001. **B.** Representative mages and replicate quantitation of RFP/GFP ratio in RPE1 RPS3-RFP-GPF cells treated with Mps1i pulse or DMSO (Ctrl), collected at 24 or 72 hours and upon Bafilomycin A_1_ (BafiloA_1_) treatment (100nM, 6h). RFP/GFP ratio was calculated as explained in Methods and visualised with a rainbow lookup table; cyan masks indicate primary nuclei; images have been obtained from the experiments shown in Fig2F; scale bars 50μm. Upper quartile, lower quartile and median of each violin plot are shown; n=3 biological replicates (30 fields of view each); Kruskal-Wallis test, followed by Dunn’s multiple comparison test: **** indicates *p*<0.0001.

### Impaired protein folding triggers autophagic removal of ribosomes in aneuploid cells

We next sought to understand the nature of the stimuli triggering ribosomal degradation in aneuploid cells. Protein homeostasis is severely challenged in aneuploid cells by compromised activities of chaperone-mediated folding(*25*, *43*). We previously found that aneuploidy activates the unfolded protein response (UPR), resulting in attenuated translation and increased protein degradation(*23*). In agreement with these observations, we found that protein levels of a panel of well-known clients of Hsp90 chaperone are reduced in aneuploid cells (**Figure S3A**). Importantly, 72 hours after induction of chromosome mis-segregation, this reduction is comparable to that obtained upon treatment of parental cells with the Hsp90 inhibitor Geldanamycin (**Figure S3A**). The impairment of chaperone-mediated folding is also reflected in the aggregation of misfolded and unfolded proteins (**Figure S3B**), detected by using the tetraphenylethene maleimide dye (TMI), that fluoresces once it reacts with free cysteine thiols that get exposed when proteins are unfolded(*44*). To determine the extent of increased protein folding demand in aneuploid cells and the impact of the consequent folding defects on ribosomal function, we inhibited chaperone activity in parental cells. Interestingly, treatments with Hsp90 inhibitors - Geldanamycin or 17-AAG - and with Hsp70 inhibitor VER-155008 led to increased degradation of ribosomes in HCT116 cells (**Figure 4A-B** and **Figure S3C**) and in RPE1 cells (**Figure S3D-E**). Importantly, inhibition of HSP90 and HSP70 had an impact specifically into ribosomal degradation and did not affect bulk autophagy, as shown by lack of processed Keima band in HCT116 and RPE1 cells expressing LDHB-Keima (**Figure S3F-G**). These data indicate that impaired chaperone activity does not trigger bulk autophagy, further suggesting that its effect is specific to the autophagic degradation of ribosomes. Thus, impaired chaperone-mediated folding is likely the main trigger of ribosome degradation. This suggests that chaperones might be limiting in aneuploid cells and their folding capacity could be saturated due to their overwhelming caused by aneuploidy. This is due to increased folding demand brought about by the aneuploid karyotypes and the lack of binding partners able to interact with the emerging polypeptide chain from the ribosome. In turn, this might lead to queuing of nascent polypeptides from the ribosomes, leading to a transient stalling in protein synthesis. To test this hypothesis, we reasoned that restoring the folding process in aneuploid cells might rescue the degradation of ribosomes. To assess this, we first induced the expression of chaperones by employing a constitutively active form of HSF1, a master regulator of chaperones(*25*). In agreement with our hypothesis, the HSF1-dependent expression of HSP27 and HSP70 in aneuploid cells significantly reduced the degradation of ribosomes via autophagy (**Figure 4C**). Likewise, transient overexpression of Hsp90 led to a substantial decrease in ribosome degradation (**Figure 4D**). Together, these results indicate that restoring protein folding in aneuploid cells prevents the degradation of ribosomes via autophagy.

**Figure 4.**
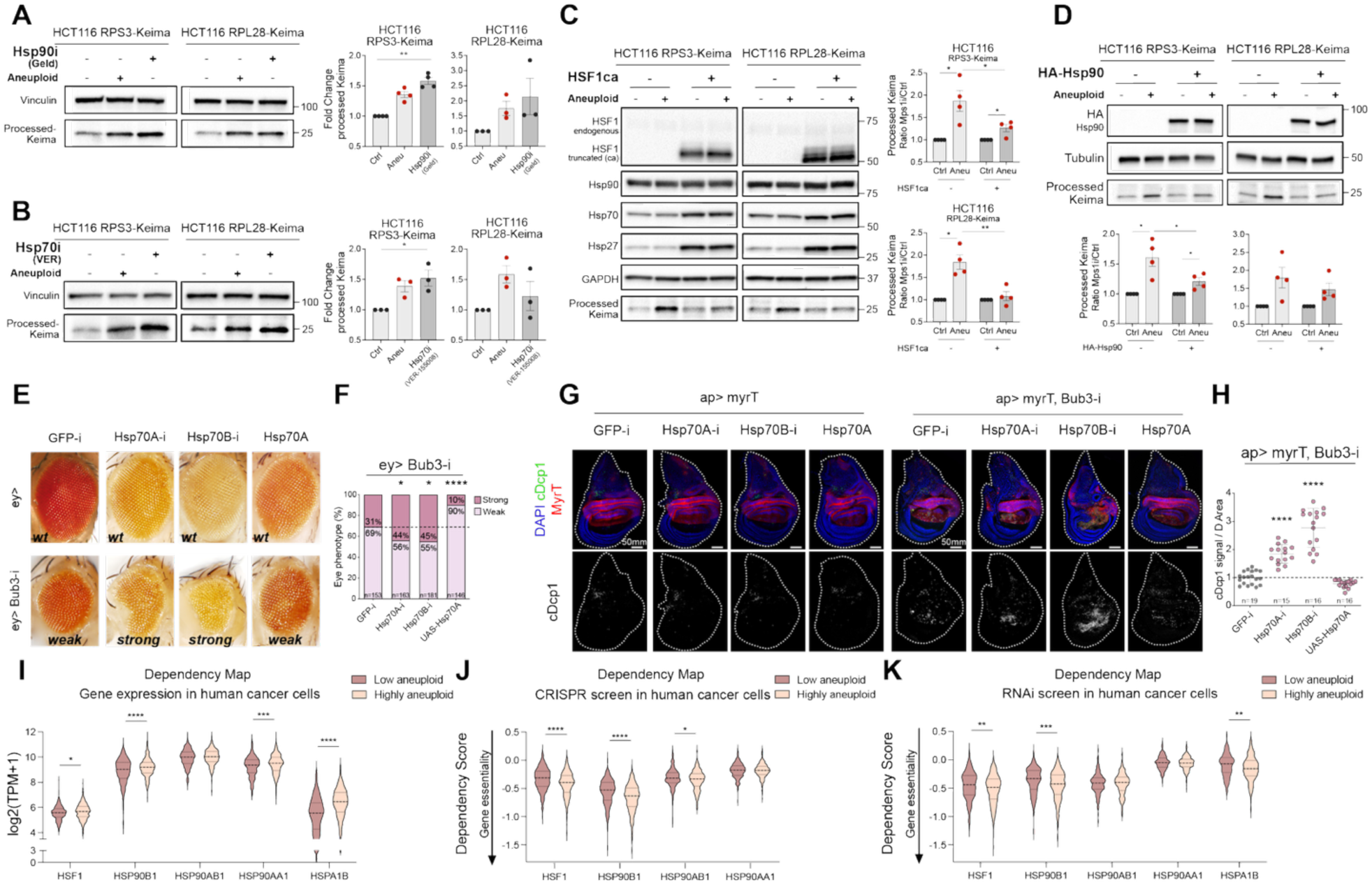
Impaired protein folding is involved in the autophagic removal of ribosomes in aneuploid cells. **A.** Representative immunoblots and replicate quantitation of processed-Keima levels in the indicated HCT116 Ribo-Keima cell lines treated for 24 hours with Mps1i, DMSO (control) or Hsp90i Geldanamycin (1μM); vinculin was used as loading control. Mean ± SEM, *n*=4 for HCT116 RPS3-Keima, *n*=3 for HCT116 RPL28-Keima; Kruskal-Wallis test, followed by Dunn’s multiple comparison test: ** indicates *p*=0.0042. **B.** Representative immunoblots and replicate quantitation of processed-Keima levels in the indicated HCT116 Ribo-Keima cell lines treated for 24 hours with Mps1i, DMSO (control) or Hsp70i VER-155008 (50μM); vinculin was used as loading control. Mean ± SEM, *n*=3; Kruskal-Wallis test, followed by Dunn’s multiple comparison test: * indicates *p*=0.0459. **C.** Representative immunoblots and replicate quantitation of processed-Keima levels in the indicated HCT116 Ribo-Keima cell lines upon HSF1ca (or empty backbone) expression and treated for 24 hours with Mps1i or DMSO (control); HSF1 as overexpression control, Hsp90, Hsp70 and Hsp27 were blotted and GAPDH was used as loading control. Mean ± SEM, *n*=4; one sample and Wilcoxon test (Mps1i *vs* respective control=1): * indicates *p*=0.0335 (HCT116 RPS3-Keima Ctrl *vs* Mps1i) or *p*=0.0475 (HCT116 RPS3-Keima HSF1ca Ctrl *vs* Mps1i) or *p*=0.0128 (HCT116 RPL28-Keima Ctrl *vs* Mps1i); unpaired Student’s t-test: * indicates *p*=0.0486 (HCT116 RPS3-Keima Mps1i *vs* HSF1ca Mps1i), ** indicates *p*=0.0065 (HCT116 RPL28-Keima Mps1 *vs* HSF1ca Mps1i). **D.** Representative immunoblots and replicate quantitation of processed-Keima levels in the indicated HCT116 Ribo-Keima cell lines upon HA-Hsp90 (or mock) overexpression and treated for 24 hours with Mps1i or DMSO (control); HA was blotted as overexpression control and tubulin was used as loading control. Mean ± SEM, *n*=4; one sample and Wilcoxon test (Mps1i *vs* respective control=1): * indicates *p*=0.0267 (HCT116 RPS3-Keima Ctrl *vs* Mps1) or *p*=0.0479 (HCT116 RPS3-Keima HA-HSP90 Ctrl *vs* Mps1); unpaired Student’s t-test: * indicates *p*=0.0485 (HCT116 RPS3-Keima Mps1 *vs* HA-Hsp90 Mps1). **E.** Representative adult eyes in which CIN was induced with an eye-specific Gal4 driving bub3-RNAi (ey>bub3-i) or control wild-type eyes (ey>), where the indicated transgenes for either the depletion of Hsp70 (Hsp70A-i / Hsp70B-i) or its overexpression (Hsp70A) were expressed. **F.** Quantification of the analysed eye phenotypes, obtained as in **(e)** and expressed as the percentage of each phenotype across the samples. Number of replicates is shown on each histogram bar; Fisher’s exact test: * indicates *p*=0.0277 (Hsp70A-i) or *p*=0.0133 (Hsp70A-i), **** indicates *p*<0.0001. **G.** Representative images of larval wing discs in which CIN was induced with an apterous-specific Gal4 driving bub3-RNAi fused with a MyrT fluorescent protein (ap>myrT, bub3-i) or control wild-type tissues (ap>myrT), where the indicated transgenes for either the depletion of Hsp70 (Hsp70A-i / Hsp70B-i) or its overexpression (Hsp70A) were expressed. Tissues have been stained with cleaved-Dcp1 (cDcp1) and DAPI (to stain DNA); MyrT signal marks the aneuploid region; scale bars 50mm. **H.** Quantification of the area positive for cDcp1 signal, obtained as in **(g)** and normalised to the area of the dorsal region (D) (where ap>myrT, bub3-i is expressed) in the indicated samples. Number of replicates is shown on graph; one-way ANOVA, followed by Dunnett’s multiple comparison test (GFP-i vs each other sample): **** indicates *p*<0.0001. **I.** mRNA expression levels (log2(TPM+1)) of the reported genes, comparing the top and bottom aneuploidy quartiles of human cancer cell lines from the CCLE. Upper quartile, lower quartile and median of each violin plot are shown; two-tailed Student’s t-test: * indicates *p*=0.0185, *** indicates *p*=0.0007, **** indicates *p*<0.0001. **J.** CRISPR screens data of the reported genes, comparing the top and bottom aneuploidy quartiles of human cancer cell lines from the CCLE. Upper quartile, lower quartile and median of each violin plot are shown; two-tailed Student’s t-test: * indicates *p*=0.0292, **** indicates *p*<0.0001. **K.** RNAi dependency data of the reported genes, comparing the top and bottom aneuploidy quartiles of human cancer cell lines from the CCLE. Upper quartile, lower quartile and median of each violin plot are shown; two-tailed Student’s t-test: ** indicates *p*=0.0061 (HSF1) or *p*=0.0062 (HSPA1B), *** indicates *p*=0.0001.

Having established the role of limiting chaperone levels in compromised proteostasis of aneuploid cells, we next set out to test whether chaperones are also limiting in an *in vivo* setting. For this, we took advantage of a *Drosophila melanogaster* model with chromosomal instability (CIN), induced by depletion of the SAC gene Bub3 in the highly proliferative epithelia of the eye and wing primordia(*21*). In this fly model, CIN-induced aneuploid cells were shown to present high levels of proteotoxic stress, and to enter apoptosis(*21*). Induction of CIN in the eye (*ey>bub3-i*) produced rough eyes with a reproducible mild reduction in size (referred to as “weak” phenotype). Interestingly, in eyes subjected to CIN, depletion of Hsp70 chaperone - with two different RNAi - further enhanced the reduction of eye size and roughness (“strong” phenotype), while, notably, Hsp70 overexpression reduced the presence of these “strong” phenotypes (**Figure 4E**, bottom panels and **Figure 4F**). Same Hsp70 modifications in wild-type eyes (*ey>*), instead, did not show any deleterious effect (**Figure 4E** upper panels). Consistently, chaperone depletion also increased the number of CIN-induced apoptotic cells in larval wing imaginal disc (**Figure 4G**, right panels and **Figure 4H**), while it had no effect on otherwise wild-type tissue (**Figure 4G**, left panels). Most interestingly, Hsp70 overexpression rescued the levels of apoptosis upon CIN (**Figure 4G**, right panels and **Figure 4H**). These data support the idea that aneuploid cells are highly sensitive to the impairment of chaperone-mediated folding *in vivo*.

Importantly, highly aneuploid human cancer cells overexpress chaperones and are more dependent on them, compared to low aneuploid cancer cells. Indeed, by interrogating the Dependency Map (DepMap), we found that, across hundreds of human cancer cell lines, the highly aneuploid ones express higher levels of HSF1, Hsp90 (HSP90B1, HSP90AB1, HSP90AA1) and Hsp70 (HSPA1B) chaperones (**Figure 4I)**, consistent with their higher folding demand. Also, highly aneuploid cancer cell lines are more dependent on those genes, as found by CRISPR (**Figure 4J)** and RNAi (**Figure 4K)** screens. These observations further highlight the essentiality of chaperones in aneuploid cells, across different models of aneuploidy.

Taken together, these findings demonstrate that aneuploid cells experience chaperone overload, which may initiate a cascade of events leading to impaired translation and ultimately triggering autophagic removal of ribosomes.

### Aneuploidy impacts global protein synthesis and translation efficiency

Ribosomes translate genetic information into proteins(*24*). Given the fact that ribosomes are actively recognized and degraded in aneuploid cells, we wanted to investigate whether and how translation is impacted in cells harbouring aneuploid karyotypes. Thus, we measured global protein synthesis rates by SunSET (Surface Sensing of Translation) assay, which consists of puromycin incorporation into newly synthesised polypeptides(*45*). We found that aneuploid populations - from both HCT116 and RPE1 cells - incorporated less puromycin than control samples, over a 30-minute incubation (**Figure 5A**), indicating that global protein synthesis rates are reduced in aneuploid cells. Protein synthesis defects accumulated over time (most likely due to a gradual build-up of aneuploidy-associated stresses, **Figure S4A**), and correlated with increasing degrees of aneuploidy (**Figure S4B**), which is in line with our recent findings(*23*). Next, to confirm the functional relationship between proteotoxic stress due to limited chaperone activities and protein synthesis attenuation, we inhibited Hsp90 activity with Geldanamycin, and we found that this treatment further decreased protein synthesis rates (**Figure 5B**). Importantly, consistent with the observation that restoring the folding process in aneuploid cells rescued ribosome degradation (**Figure 4C, D**), we found that HSF1-dependent expression of HSP27 and HSP70 significantly restored protein synthesis (**Figure 5C**).

**Figure 5.**
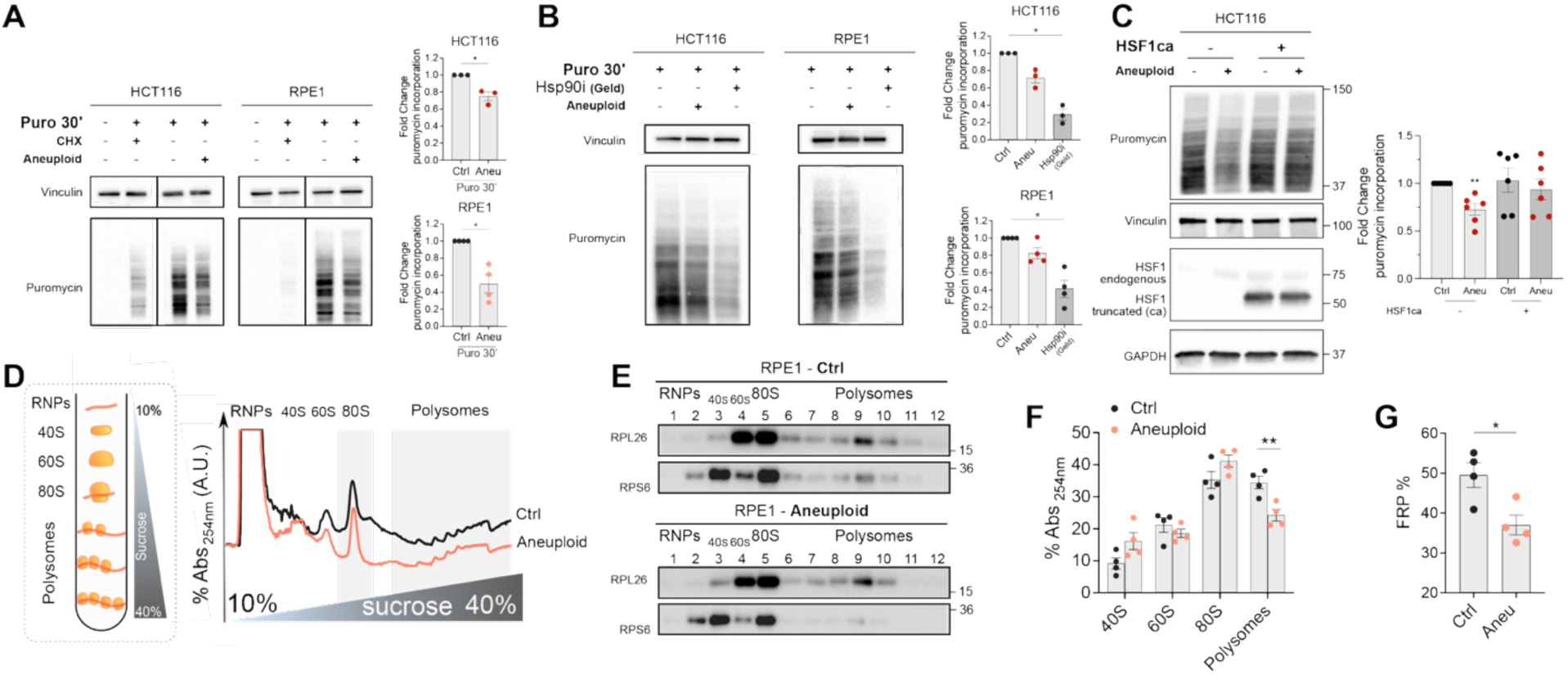
Aneuploidy impacts global protein synthesis and translation efficiency. **A.** Representative immunoblots and replicate quantitation of puromycin incorporation (10μg/mL over 30 minutes) into the indicated cell lines, treated with Mps1i pulse or DMSO (control) and collected at 24 hours (HCT116) or 72 hours (RPE1); treatment with cycloheximide (CHX) (10μg/mL, 6h) was used as a negative control; vinculin was used as loading control for puromycin smear quantitation. Mean ± SEM, *n*=3 for HCT116, *n*=4 for RPE1; one sample and Wilcoxon test (Mps1i *vs* respective control=1): * indicates *p*=0.0382 (HCT116) or *p*=0.0170 (RPE1). **B.** Representative immunoblots and replicate quantitation of puromycin incorporation (10μg/mL over 30 minutes) into the indicated cell lines treated for 24 hours with Mps1i, DMSO (control) or Hsp90i Geldanamycin (1μM); vinculin was used as loading control for puromycin smear quantitation. Mean ± SEM, *n*=3 for HCT116, *n*=4 for RPE1; Kruskal-Wallis test, followed by Dunn’s multiple comparison test: * indicates *p*=0.0190 (HCT116) or *p*=0.0156 (RPE1). **C.** Representative immunoblots and replicate quantitation of puromycin incorporation (10μg/mL over 30 minutes) into the indicated HCT116 cell line, upon HSF1ca (or empty backbone) expression and treated for 24 hours with Mps1i or DMSO (control); vinculin and GAPDH were used as loading control. Mean ± SEM, *n*=6; one sample and Wilcoxon test. ** indicates *p*=0.0063. **D.** Schematic showing polysome profiling (left) and representative sucrose gradient absorbance at 254nm profiles (right) obtained from the cytoplasmic lysates of RPE1 cells treated with Mps1i pulse or DMSO (Ctrl) and analysed at 72 hours. **E.** Representative immunoblots for the co-sedimentation profiles of two ribosomal subunit markers (RPL26 and RPS6) in the sucrose gradient fractions obtained as in **(D)**. **F.** Quantitation of the relative absorbance (at 254nm) distribution of each component in Mps1i-and DMSO (Ctrl)-treated samples, calculated from the profiles obtained as in **(D)**. Mean ± SD; *n*=4; unpaired Student’s t-test: ** indicates *p*=0.0081. **G.** Comparison between the fraction of ribosomes in polysomes (FRP%) in Mps1i- and DMSO (Ctrl)-treated samples, calculated from the profiles obtained as in **(D)**. Mean ± SD; *n*=4; unpaired Student’s t-test: * indicates *p*=0.0201.

To provide additional insight on translational alterations, we performed polysome profiling, which provides information on the mRNA engagement with the translational machinery and the relative distribution of ribosomes in a cytoplasmic lysate (**Figure 5D**). Analysis of profiles (**Figure 5D**) as well as the resulting co-sedimentation blots (**Figure 5E**) indicates that while the ribosomal subunits and the ribosome (80S) did not change their relative distribution, the polysomal fraction was significantly decreased in aneuploid samples (**Figure 5F**). Next, we calculated the fraction of ribosomes loaded on <u>p</u>olysomes (FRP) and found this to be strongly reduced in aneuploid cells (**Figure 5G**), confirming that their translation efficiency is decreased.

Overall, these data demonstrate that aneuploid cells experience reduced translation efficiency, likely due to folding stress, ultimately leading to the removal of translationally impaired ribosomes.

### The E3-ligase ZNF598 mediates autophagic removal of ribosomes in aneuploid cells

Defects in ribosome functioning are actively recognized by the Ribosome-associated Protein Quality Control (RQC) system, a cellular surveillance mechanism that identifies stalled ribosomes, mediates their dissociation, and promotes their degradation(*32–34*). Given the translational and folding defects observed in aneuploid cells, we decided to test whether the RQC machinery contributes to managing aneuploidy-induced proteotoxic stress. Specifically, we focused on the E3-ligase ZNF598(*46*), which is responsible for recognizing defective ribosomes and subsequently triggering the RQC(*34*). To explore ZNF598’s involvement in ribosome degradation in aneuploid cells, we depleted ZNF598 and induced chromosome mis-segregation. We found that loss of ZNF598 results in a significant decrease in the autophagic degradation of ribosomes, observed 24 hours after chromosome mis-segregation (**Figure 6A, B**). Since ZNF598 functions through the specific ubiquitylation of substrates, namely proteins of the small ribosomal subunit(*46*), we examined this process in more detail. We immunoprecipitated ubiquitylated proteins from both aneuploid and control samples and performed blotting for the ribosomal protein RPS3 in the presence and absence of ZNF598. We found RPS3 to be enriched in the ubiquitylated pool in aneuploid cells, and this enrichment was reduced upon ZNF598 depletion (**Figure S5A**), confirming that ribosome degradation depends on ZNF598-mediated ubiquitylation. These findings suggest that autophagy-driven removal of ribosomes occurs in aneuploid cells upon recognition of their translationally impaired state, a process mediated by ZNF598.

**Figure 6.**
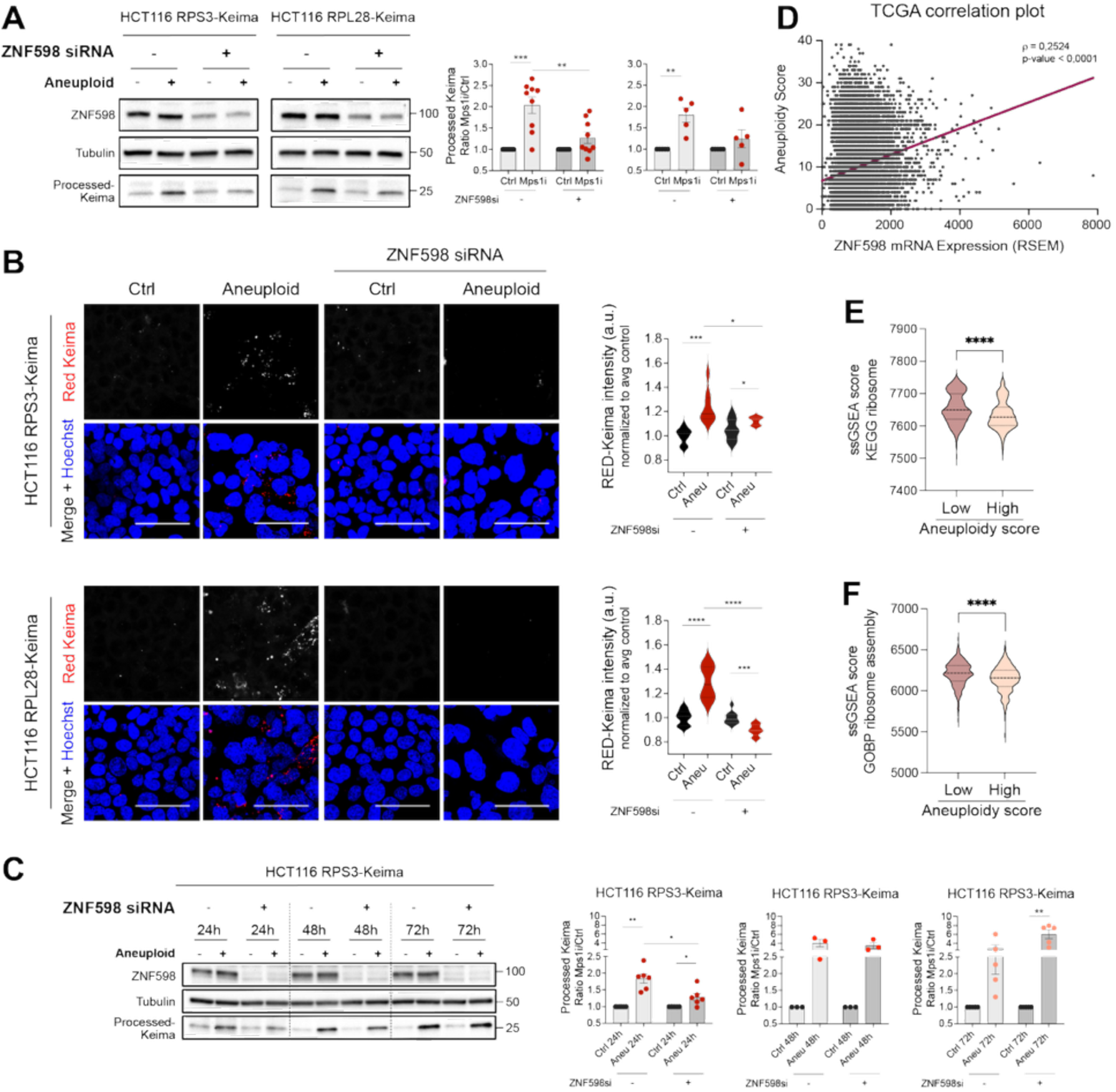
The E3-ligase ZNF598 mediates the autophagic removal of ribosomes in aneuploid cells. **A.** Representative immunoblots and processed-Keima quantitation of the indicated HCT116 Ribo-Keima cell lines upon ZNF598 siRNA (or non-targeting siRNA) and treated for 24 hours with Mps1i or DMSO (control); ZNF598 was blotted as knock-down control and tubulin used as loading control. Mean ± SEM, *n*=9 for HCT116 RPS3-Keima, *n*=5 for HCT116 RPL28-Keima; one sample and Wilcoxon test (Mps1i *vs* respective control=1): ** indicates *p*=0.0096, *** indicates *p*=0.0006; unpaired Student’s t-test (Mps1i *vs* ZNF598siRNA Mps1i): ** indicates *p*=0.0049. **B.** Representative live-cell images and replicate quantitation of indicated HCT116 Ribo-Keima cells lines upon ZNF598 siRNA (or non-targeting siRNA) and treated for 24 hours with Mps1i pulse or DMSO (Ctrl). Red-Keima intensity is proportional to ribosome degradation; Hoechst was used to stain DNA; scale bars, 50μm. Upper quartile, lower quartile and median of each violin plot are shown, *n*=9 fields of view; unpaired Student’s t-test: * indicates *p*=0.0272 (HCT116 RPS3-Keima Mps1i *vs* ZNF598si Mps1i) or *p*=0.0359 (HCT116 RPS3-Keima ZNF598si Ctrl *vs* Mps1i), *** indicates *p*<0.0004 (HCT116 RPS3-Keima Ctrl *vs* Mps1i) or *p*=0.0005 (HCT116 RPL28-Keima ZNF598si Ctrl *vs* Mps1i), **** indicates *p*<0.0001. **C.** Representative immunoblots and processed-Keima quantitation of HCT116 RPS3-Keima cells upon ZNF598 siRNA (or non-targeting siRNA), treated with Mps1i pulse or DMSO (control) and collected at 24, 48 or 72 hours; ZNF598 was blotted as knock-down control and tubulin was used as loading control. Mean ± SEM, *n*=6 for 24-hour samples, *n*=3 for 48-hour samples, *n*=5 for 72-hour samples; one sample and Wilcoxon test (each Mps1i *vs* respective control=1): * indicates *p*=0.0449, ** indicates *p*=0.0017 (Ctrl vs Mps1i 24h) or *p*=0.0072 (ZNF598si Ctrl vs Mps1i 72h), unpaired Student’s t-test (each Mps1i *vs* ZNF598siRNA Mps1i): * indicates *p*=0.0101. **D.** Pan-cancer correlation plot between ZNF598 (RSEM) mRNA expression levels and the aneuploidy scores from TCGA primary tumour samples. The trend line for Spearman’s correlation is in purple: ρ=0,2524, *p*<0.0001. **E, F.** Association between ssGSEA score for GO Biological Process ribosome assembly (E) or KEGG ribosome signature (F) and aneuploidy scores, comparing the top and bottom aneuploidy quartiles of human cancer cell lines from the CCLE. Upper quartile, lower quartile and median of each violin plot are shown; unpaired Student’s t-test: **** indicates *p*<0.0001.

Ribosome recognition is primarily achieved through ZNF598, which detects stalled ribosomes by recognizing collided di-ribosomes, a structural hallmark of translational impairment (*46*). This recognition allows ZNF598 to target only defective ribosomes for degradation, maintaining translational quality control. Importantly, it has been proposed that under conditions of strong autophagic activation—such as during nutrient starvation(*39*)—ribosomes can be degraded via a non-specific, bulk autophagy process, as part of the cell’s adaptive response to stress. This autophagic process facilitates nutrient recycling and helps maintain cellular homeostasis when resources are scarce(*47*). Because of this, we hypothesized that the autophagic removal of ribosomes in aneuploid cells might depend on ZNF598 when ribosomes are the primary cargo and no other major autophagic substrates are present— a condition we observed approximately 24 hours after chromosome mis-segregation (**Figure 1D, E** and **Figure S1A-D**). Conversely, when multiple cargos are targeted for degradation through autophagy—as we found occurring at later time points after chromosome mis-segregation (e.g., 48 and 72 hours; **Figure S1A–D**)—ribophagy may become a bystander process, rendering ZNF598 dispensable. In agreement with this idea, we found that autophagic degradation of ribosomes 48 and 72 hours after chromosome mis-segregation was not impacted upon ZNF598 depletion (**Figure 6C** and **Figure S5B-C**). Thus, we speculate that ribosomes may initially be degraded through a process primarily mediated by ZNF598—immediately following chromosome missegregation—when the cellular stresses induced by aneuploidy are still manageable and stalled ribosomes constitute the main autophagic cargo. During this early phase, aneuploid cells undergo proteostasis disruption due to imbalances in protein stoichiometry caused by chromosome number imbalances. In this stage, the cell likely depends on precise mechanisms to eliminate translationally defective ribosomes, aiming to maintain proteostasis. However, as the cellular stresses accumulate and become more severe, the autophagic response shifts to a bulk autophagy mode (**Figure S1A-D**). In this later stage, autophagy targets multiple cellular components—including ribosomes, ER, and other organelles—for degradation, as part of an overarching effort to recycle nutrients and promote survival under stress conditions(*40*).

Since ribosome surveillance mediated by ZNF598 may serve as a crucial quality control mechanism in aneuploid cells – helping them managing proteostasis disruption – and given that the majority of cancers harbour aneuploid karyotypes(*2*), we sought to analyse ZNF598 expression in tumour samples. To do this, we analysed data from the TCGA (The Cancer Genome Atlas) database, examining the association between ZNF598 mRNA levels and aneuploidy scores across more than 11000 primary tumour samples. This pan-cancer analysis revealed a significant positive correlation between ZNF598 expression and tumour aneuploidy scores (ρ=0.25, p<0.0001) (**Figure 6D**). This suggests that aneuploid tumors might upregulate ZNF598 as part of a cellular stress response aimed at maintaining proteostasis through ribosome degradation. To further validate this, we analyzed gene expression data from the CCLE (Cancer Cell Line Encyclopedia), focusing on cell lines with high versus low levels of aneuploidy. We observed that highly aneuploid cancer cell lines tend to downregulate pathways related to ribosome biogenesis, protein synthesis, and translation compared to pseudo-diploid lines (**Figure 6E, F**). This supports the hypothesis that these cells actively modulate their translational machinery in response to aneuploidy. Additionally, the analysis of ribosomal and translational gene signatures revealed an inverse relationship with ZNF598 expression (**Figures S6A–B**). Specifically, high ZNF598 levels were associated with a significant decrease in the expression of genes involved in ribosome biogenesis, protein synthesis, and the overall translational machinery. We hypothesize that this reflects a broader adaptive response whereby cancer cells attenuate cytotoxic stresses at the levels of the ribosome, thereby aiming at improving proteostasis. This pattern also suggests that upregulation of ZNF598 may serve as a cellular strategy to mitigate the proteotoxic and folding stresses induced by aneuploidy. In this context, ZNF598 may facilitate regulation of key transcripts necessary for continued proliferation and survival. One prominent candidate is c-Myc, which is involved in cancer cell proliferation, whose translation has recently been shown to be regulated by ZNF598 in glioblastoma(*48*). Our data support this connection. Indeed, we observed a positive correlation between ZNF598 and c-Myc expression across cancer cell lines (**Figure S6C**). This suggests that cancer cells, especially those with extensive chromosomal imbalances, undergo a broad attenuation of the cellular stresses associated with aneuploidy—such as proteotoxic and folding stress, among others(*23*, *49*, *50*)—and that ZNF598 may facilitate this adaptation by regulating the translation of critical oncogenic transcripts. This enables ZNF598 to target defective ribosomes for degradation and preserve translation.

Altogether, our data indicate that ZNF598 plays a key role in orchestrating the degradation of translationally impaired ribosomes in aneuploid cells. Additionally, ZNF598 may act as a crucial regulator balancing proteostasis and proliferation in highly aneuploid cancers.

## DISCUSSION

Proteotoxic stress is a hallmark of aneuploidy, primarily driven by gene copy number alterations that disrupt protein balance and lead to the accumulation of misfolded proteins. This overloads the cell’s proteostasis machinery, including chaperones, the ubiquitin-proteasome system, and autophagy(*4*, *23*, *26*, *49*, *51*, *52*). As a result of this, aneuploid cells are especially sensitive to additional stresses exacerbating proteotoxic stress, because of their already-compromised folding machinery(*25*, *43*). Recent studies have shed light on the pathways involving the unfolded protein response and selective autophagy in aneuploid cells(*13*, *23*, *49*). However, a comprehensive understanding of the mechanisms activated during the onset of proteotoxicity, as well as the fundamental processes by which cells detect and effectively resolve proteotoxic stress, remains unclear. Gaining a better understanding of these processes could help identify vulnerabilities in aneuploid cancers, which might lead to the development of novel therapies in the future.

Here, we present a molecular dissection of the events taking place after chromosome mis-segregation in response to the disruption of protein homeostasis. By combining multiple reporters to tag autophagic cargo with protein-based assays and live-cell imaging experiments, mass spectrometry and correlative light-electron microscopy, we demonstrate that ribosomes are actively removed in aneuploid cells (**Figure 1-3**). We found that this process begins with impaired chaperone-mediated folding, eventually leading to reduced translation (**Figure 4-5**). Our in vivo experiments further support the notion that aneuploid cells are particularly vulnerable to disruptions in chaperone-mediated folding and proteostasis. Notably, highly aneuploid cancer cells display an increased dependence on chaperones, along with elevated expression of related genes (**Figure 4**), suggesting that protein folding becomes a limiting factor in cells carrying chromosomal imbalances.

Our results further suggest that translationally defective ribosomes are detected by the E3 ligase ZNF598, which tags these ribosomes for degradation, thus helping cells managing proteostasis disruption during early stress. Moreover, we also demonstrate that as stress worsens, autophagy shifts to a more general degradation mechanism, targeting a wide range of cellular components including ribosomes, endoplasmic reticulum, mitochondria, and other organelles(*49*, *53*). This broad activity aims to remove damaged or dysfunctional structures and provide energy for cell survival. As a result, at this stage, ribophagy becomes a bystander process in aneuploid cells and ZNF598’s activity is dispensable (**Figure 6, 7**).

**Figure 7.**
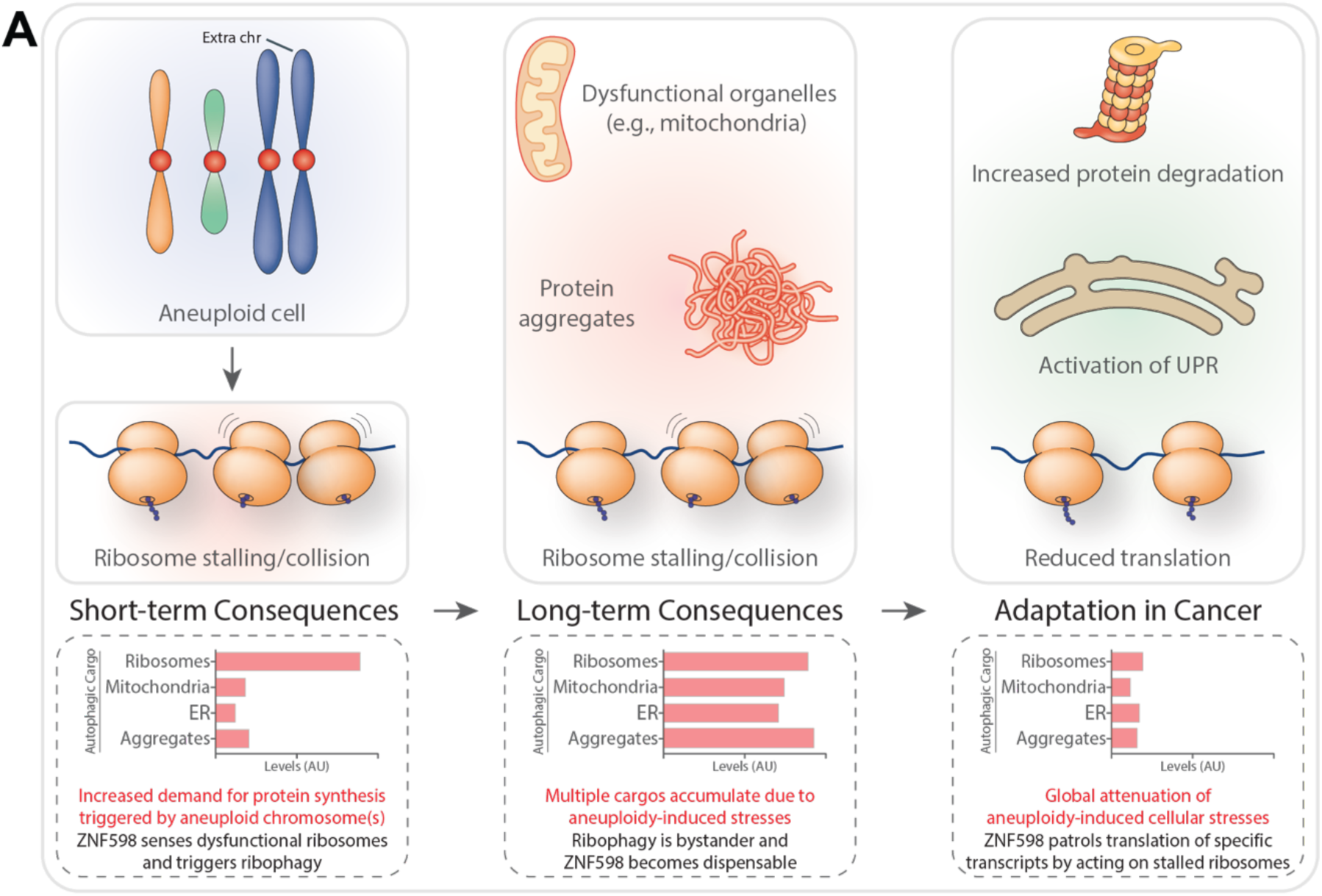
Model for the consequences of aneuploidy on proteostasis and how cancer cells adapt to these stresses. **A.** Schematic model illustrating the cellular events occurring in aneuploid cells upon proteostasis disruption. Immediately after chromosome mis-segregation, there is an increased demand for protein synthesis due to the presence of aneuploid chromosomes. During this phase, ZNF598 detects dysfunctional ribosomes and activates ribophagy to remove them. As aneuploidy-induced stresses cause multiple cargos to accumulate, ribophagy shifts to a more non-specific, bystander role, making ZNF598 dispensable. Over time, as cancer cells adapt, they undergo a broad reduction of the stresses induced by aneuploidy, including global reduction of translation. Under these conditions, ZNF598 regulates the translation of specific transcripts by acting on stalled ribosomes. See text for more details.

The vast majority of tumors are aneuploid(*1–4*). Given the negative impact of aneuploidy on cell physiology, it is likely that cancer cells have developed mechanisms to adapt to these stresses. Supporting this, recent studies have shown that during cancer progression, there is a widespread reduction in aneuploidy-induced cellular stresses—such as increased degradation of mRNA and proteins, along with decreased replication stress and genomic instability(*12*, *23*, *49*, *54*). Also, it has been suggested that protein buffering and the maintenance of proper protein complex stoichiometries are common features in cancer cells and are crucial for supporting sustained cancer cell proliferation(*55*, *56*). Our data strongly support these concepts and suggest that ZNF598, along with other components of the RQC, may regulate key transcripts essential for tumor proliferation and survival (**Figure 7A**). Recent studies in glioblastoma have shown that ZNF598 can modulate c-Myc translation(*48*), and our findings indicate this regulatory mechanism may be conserved across multiple cancer types. This adds an additional layer of control at the translational level during adaptation to aneuploidy in cancer (**Figure 7A**).

Therefore, we propose that the reduction in ribosome biogenesis, combined with increased ZNF598 expression, reflects a widespread suppression of aneuploidy-induced stresses at the ribosome level, unveiling a novel vulnerability that could be targeted in cancer therapy. Future research in this area is expected to deepen our understanding of how aneuploid cancer cells adapt to proteotoxicity. Since several compounds that interfere with ribosome function are currently in clinical trials and show preferential toxicity to cancer cells(*57*, *58*), targeting ribosome quality control offers a promising strategy. The notion that highly aneuploid cancers might be more sensitive to these drugs is particularly intriguing and could pave the way for new approaches in predicting therapy response and developing prognostic biomarkers.

### Materials and Methods Cell culture conditions

hTERT RPE-1 cells, hTERT RPE-1 cells expressing RPS3-Keima or RPL28-Keima (generated in house), hTERT RPE-1 cells expressing LDHB-Keima, RAMP4-Keima or MT-Keima (all generated in house), hTERT RPE-1 cells expressing RPS3-RFP-GFP (generated in house) and HEK-293T were cultured in Dulbecco’s modified Eagle’s High Glucose medium (DME/HIGH with stable L-Glutamine and Sodium Pyruvate) supplied with 10% FBS (South America origin) and 100U/mL Penicillin/Streptomycin. HCT116 cells, HCT116 cells expressing RPS3-Keima or RPL28-Keima (kindly provided by Prof. Wade Harper), HCT116 cells expressing LDHB-Keima, RAMP4-Keima or MT-Keima (all generated in house) and HCT116 cells expressing RPS3-RFP-GFP (generated in house) were cultured in Mc Coy’s 5A medium supplied with 10% FBS (South America origin), 2mM L-Glutamine and 100U/mL Penicillin/Streptomycin. All cell lines were previously tested free of mycoplasma contamination with MycoAlert kit (Lonza) according to manufacturer’s instructions, followed by PCR confirmation with previously reported oligonucleotides of different mycoplasma species(*59*). All cell lines were grown at 37°C with 5% CO_2_ in a humidified incubator.

### Cell treatments

To induce random aneuploidy, cells were pulsed with 500nM (or otherwise indicated) Mps1 inhibitor (Mps1i) reversine (Cayman Chemical) for 24 hours, while, to generate the relative controls, cells were pulsed with the vehicle dimethyl sulfoxide (DMSO) for 24 hours. Sample analyses were performed either immediately after the pulse (“24h” samples) or after 1 o 2 cell cycles (“48h” or “72h” samples, in these cases Mps1i/DMSO were washed-out after the pulse), or at different time-points as reported in figure legends. To evaluate the effects of the increasing aneuploidy degree on ribosome degradation or on global protein synthesis rate, cells were treated for 24 hours with increasing concentrations of Mps1i reversine (125nM, 250nM or 500nM).

To modulate autophagy, cells were treated either with SAR405 (1μM, Cayman Chemical), Bafilomycin A1 (BafiloA1, 100nM, Sigma-Aldrich) or Torin1 (250nM, Tocris), as indicated in figure legends.

To inhibit Hsp90 chaperone family, cells were treated either with Geldanamycin (Geld, 1μM, Tocris) or 17-allylamino-17-demethoxy-geldanamycin (17-AAG, 1μM, Tocris), as indicated in figure legends. To inhibit Hsp70 chaperone family, cells were treated with VER-155008 (VER, 50μM, Sigma-Aldrich), as indicated in figure legends. To induce the aggregation of misfolded proteins, cells were treated 24 hours with L-azetidine-2-carboxylic acid (AZC, 10mM, Sigma-Aldrich).

To inhibit translation elongation, cells were treated with Cycloheximide (CHX, 10μg/mL, Sigma-Aldrich) as indicated in the corresponding figure legend.

### Stable cell line generation

To generate hTERT RPE-1 or HCT116 cells stably expressing RPS3-Keima, RPL28-Keima, LDHB-Keima, RAMP4-Keima, MT-Keima or RPS3-RFP-GFP, lentivirus particles were produced by HEK-293T transfection with 10μg plasmid DNA of interest (see **Table 1**)/viral packaging plasmids/CaCl_2_/HBS mixture. Target cells (either hTERT RPE-1 or HCT116) where then infected with viral particles produced in HEK-293T and selected with puromycin (Sigma-Aldrich, 10μg/mL for hTERT RPE-1 or 3μg/mL for HCT116) or blasticidin (Santa Cruz Biotechnology, 5μg/mL for hTERT RPE-1 or 6μg/mL for HCT116), depending on the specific resistance of each plasmid (see **Table 1**).

**Table 1:** List of the plasmids used to generate stable cell lines.

| Construct | Vector name | Source | Bacterial growth strain | Bacterial resistance | Eukaryotic resistance |
| --- | --- | --- | --- | --- | --- |
| LDHB-Keima | pHAGE LDHB-mKEIMA | Prof. Wade Harper | Stbl3 | Ampicillin | Puromycin |
| MT-Keima | pHAGE MT-mKEIMA | Prof. Wade Harper | Stbl3 | Ampicillin | Puromycin |
| RAMP4-Keima | pDEST mKEIMA-RAMP4 | Prof. Carmine Settembre | Stbl3 | Ampicillin | Blasticidin |
| RPL28-Keima | pHAGE RPL28 mKEIMA | Prof. Wade Harper | Stbl3 | Ampicillin | Puromycin |
| RPS3-Keima | pHAGE RPS3-mKEIMA | Prof. Wade Harper | Stbl3 | Ampicillin | Puromycin |
| RPS3-RFP-GFP | pHAGE RSP3 RFP-GFP | Prof. Carmine Settembre | Stbl3 | Ampicillin | Puromycin |

### Transient construct overexpression

To transiently transfect HCT116 RPS3-Keima or HCT116 RPL28-Keima cells, Lipofectamine 3000 transfection reagent (Invitrogen) was used by optimising the manufacturer’s instructions, as follow. Cells were seeded at 30-40% confluence in a six-well plate and, 24 hours later, cell were transfected with 6μL Lipofectamine 3000 transfection reagent diluted in 150μL Opti-MEM reduced serum medium (Gibco) and 2μg HSF1ca (as previously reported(*25*), kindly provided by Prof. Zuzana Storchová) or 2μg HA-Hsp90 (or the corresponding empty vector) (see **Table 2** for construct source and information) diluted in 150μL Opti-MEM and P3000 reagent (2μL/μg DNA). Then, the DNA master mix was added to the diluted Lipofectamine 3000 mix and incubated for 10 minutes at room temperature; afterwards, 300μL of DNA-lipid complexes were added to target cells and, 8 hours later, medium was changed to avoid Lipofectamine toxicity. To analyse the effects of HSF1ca or Hsp90 overexpression on aneuploid cells, 500nM Mps1i (or vehicle DMSO) was added for 24 hours before sample collection (72h after transfection).

**Table 2:** List of the constructs used to transiently transfect HCT116 RPS3-Keima or HCT116 RPL28-Keima cell lines.

| Construct | Vector name | Source | Cat. # | Bacterial growth strain | Bacterial resistance |
| --- | --- | --- | --- | --- | --- |
| Empty pcDNA 3.1 | pcDNA3.1 | Prof. Zuzana Storchová | - | DH5α | Ampicillin |
| HA-Hsp90 | pcDNA3/HA-Hsp90 | Addgene | 22487 | DH5α | Ampicillin |
| HSF1ca | pcDNA3.1/HSF1ca | Prof. Zuzana Storchová | - | DH5α | Ampicillin |

### RNA interference

To transiently knock-down the targets of interest in HCT116 RPS3-Keima or HCT116 RPL28-Keima cells, Lipofectamine RNAiMAX transfection reagent (Invitrogen) was used by optimising the manufacturer’s instructions as follow. Cells were plated at 30% confluence in a six-well plate and, 24 hours later, cells were transfected with 9μL Lipofectamine RNAiMAX transfection reagent diluted in 150μL Opti-MEM and 3μL of MAD2 siRNA or BUB1 siRNA (20μM stock) or 3μL of ZNF598 siRNA (10μM stock) (or the corresponding non-targeting siRNA controls) (see **Table 3** for siRNA source and information) diluted in 150μL Opti-MEM. Then, the diluted siRNA mix was added to the diluted Lipofectamine RNAiMAX mix and incubated for 15 minutes at room temperature; afterwards, 250μL of siRNA-lipid complexes were added to target cells and, 8 hours later, medium was changed. To analyse the effects of MAD2 or BUB1 knock-down, samples were collected 72h after transfection. To analyse the effects of ZNF598 knock-down on control or aneuploid cells, 500nM Mps1i (or vehicle DMSO) was added for 24 hours (or otherwise indicated in figure legends) before sample collection (72h after transfection).

**Table 3:**
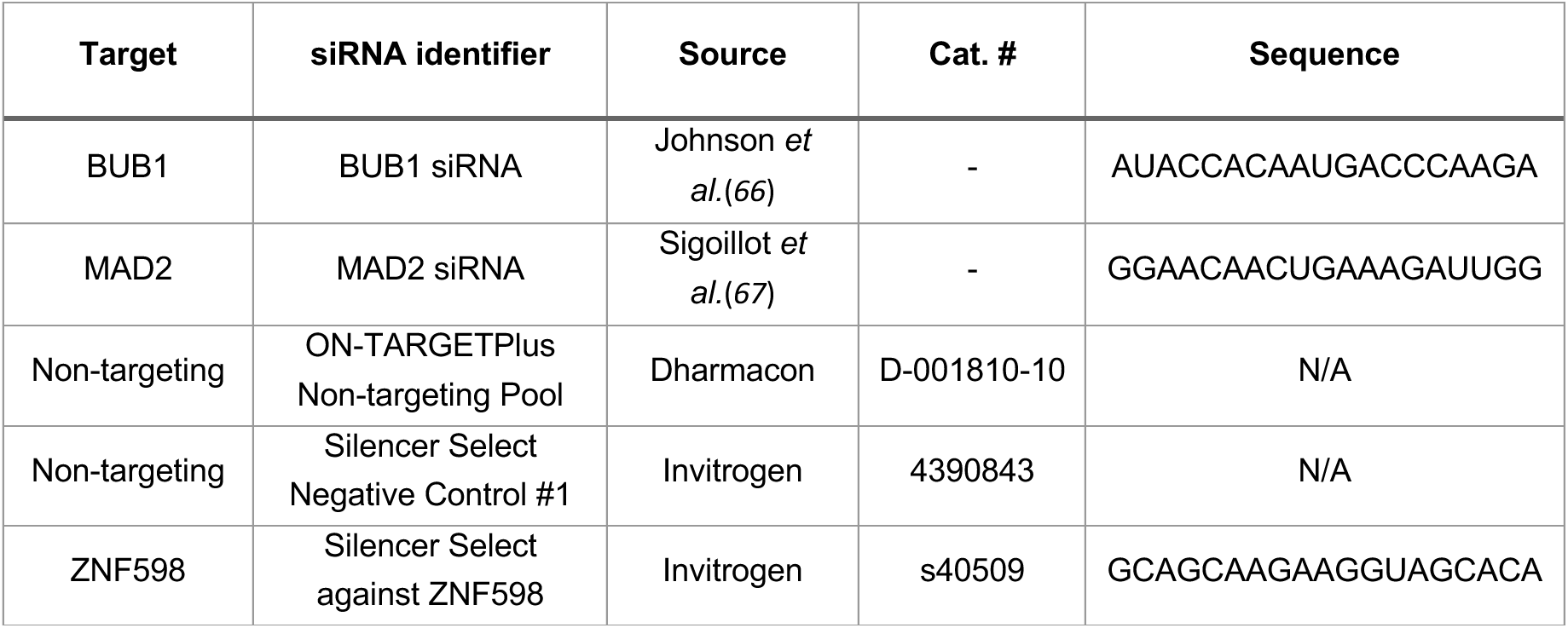
List of the siRNA sequences used to transiently knock-down the indicated targets in HCT116 RPS3-Keima or HCT116 RPL28-Keima cell lines.

### Western blot analysis

To detect proteins by Western blot, cells were lysed directly in the plate with cold RIPA Buffer (Cell Signaling Technology) supplemented with protease inhibitor cocktail (Millipore) and phosphatase inhibitor cocktail (Roche) and incubated for 10 minutes on ice. Whole-cell lysates were quantified with Bradford method-based colorimetric assay (Bio-Rad) and 30µg lysate were resolved on 4-20% Criterion TGX Stain-Free precast gels (Bio-Rad) or 4–15% Mini-PROTEAN TGX Stain-Free precast gels (Bio-Rad). Nitrocellulose membranes (Bio-Rad) with transferred proteins were blocked for 30 minutes with either 5% bovine serum albumin (BSA) or 5% non-fat milk (AppliChem GmbH) in 1XTBS+0,1% TWEEN-20 (TBS-T) at room temperature and incubated overnight at 4°C with primary antibodies diluted in 5% BSA or 5% non-fat milk in TBS-T (according to antibody datasheets). Following three washes in TBS-T, membranes were incubated 30 minutes at room temperature with secondary HRP-conjugated antibodies (anti-mouse #P0447 or anti-rabbit #P0448, Agilent) diluted 1:10000 in TBS-T, and then washed three times in TBS-T. Clarity Western ECL Substrate (Bio-Rad) was used for signal detection, images were captured on a ChemiDoc XRS + (Bio-rad) and protein band intensities were quantified with ImageLab Software (Bio-Rad) “Lane and Bands” and “Quantity tool” tools. Smear intensities from anti-puromycin immunoblots were quantified with ImageLab “Volume tool”. Each band/smear value was firstly normalised on the respective loading control value, then a second normalisation was performed on the relative control sample (DMSO-treated sample, Ctrl). The following primary antibodies were used: BUB1 (#ab54893, Abcam), EGFR (in-house polyclonal ab against human EGFR aa 1172–1186, kindly provided by Prof. Di Fiore), GAPDH (#sc-32233, Santa Cruz Biotechnology), HA (#MMS-101P, BioLegend), HSF1 (#4356S, Cell Signaling Technology), Hsp27 (#ADI-SPA-800-D, Enzo Life Science), Hsp70/72 (#ADI-SPA-810-D, Enzo Life Science), Hsp90 (#4877S, Cell Signaling Technology), Keima-red (#M182-3M, MBL), MAD2 (#A300-301A, Bethyl Laboratories), MET (#sc-162, Santa Cruz Biotechnology), mTOR (#2983S, Cell Signaling Technology), PDGFRβ (#3169S, Cell Signaling Technology), Puromycin clone 12D10 (#MABE343, Merck), Tubulin (#T9026, Sigma-Aldrich), Vinculin (#V9131, Sigma-Aldrich), ZNF598 (#ab241092, Abcam).

### Puromycin incorporation and WB-SunSET assay

To evaluate protein synthesis rates with SunSET (Surface Sensing of Translation) assay(*45*), cells were plated in order to be 70-80% confluent at the time of the assay, and the same confluence was maintained among the samples. 24 hours (HCT116) or 72 hours (RPE1) after aneuploidy induction with 500nM Mps1i pulse (or otherwise reported in figure legend) (or DMSO for controls) or 6 hours after Cycloheximide treatment, puromycin (Puromycin Dihydrochloride, Sigma-Aldrich) was diluted in cell culture medium to 10μg/mL final concentration and cells were incubated for 30 minutes at 37°C. Then, samples were lysed directly in the plates with cold RIPA buffer supplemented with protease and phosphatase inhibitors, before proceeding with Western blot analysis as described above, resolving 30µg of each sample.

### Immunoprecipitation assay

For the ubiquitylated protein immunoprecipitation experiment, HCT116 cells were seeded and transfected with ZNF598 siRNA, as described in “RNA interference” protocol paragraph. 24 hours after 500nM Mps1i treatment, 3*10^6 cells were collected and lysed on ice with 300μL of the following lysis buffer: 50mM Tris-Hcl (pH 7.5), 150mM NaCl, 0,05% NP-40, protease inhibitor cocktail (Millipore) and phosphatase inhibitor cocktail (Roche); lysates were centrifuged at 4°C for 10 minutes at 12000xg and Input samples were saved. 300μL of cell lysates were incubated with 20μL slurry anti-Multi Ubiquitin mAb-Agarose beads (MBL), for 2 hours, at 4 °C, with gentle agitation on wheel. After collection of supernatant fractions (Unbound samples), beads were washed three times with cold lysis buffer (10 seconds, 2500xg centrifugation). 10μL of 4x Laemmli’s sample buffer (supplemented with 100mM DTT) was added to the beads for SDS–PAGE and immunoblotting analysis, performed together with 20μg of Input and Unbound samples, as controls. Samples were boiled at 96°C before being resolved on a 4-20% Criterion TGX Stain-Free precast gel (Bio-Rad). Nitrocellulose membranes (Bio-Rad) with transferred proteins were incubated overnight at 4°C with FK2 (#04-263, Merck) and RPS3 (Cell Signaling Technology) primary antibodies and 30 minutes at room temperature with secondary HRP-conjugated antibodies (anti-mouse #P0447 or anti-rabbit #P0448, Agilent). RPS3 bands were quantified with ImageLab Software (Bio-Rad), values from IP samples were firstly normalised on the respective Input samples and, then, on the relative control sample (DMSO-treated sample, Ctrl).

### Quantitative real-time PCR analysis

To quantify ZNF598 expression levels, cells were pulsed with 500nM Mps1i (or DMSO) and processed at 24, 48 or 72 hours. RNA was extracted using RNeasy Plus Mini Kit (QIAGEN), following manufacturer’s instructions, and 500ng RNA were retro-transcribed in 20μL DEPC-treated water (Invitrogen) using OneScript Plus cDNA Synthesis Kit (abm), according to manufacturer’s protocol. cDNA was diluted to 6,25ng/μL and 2μL were used to perform quantitative real-time PCR, carried out using Fast SYBR Green Master Mix (ThermoFisher Scientific). Quantitative Real-Time PCR reactions were run on a CFX 96 Real-Time PCR system (Bio-Rad). mRNA relative expression levels were calculated with the 2^-ΛΛCt^ method and were shown on the graphs as fold changes: data normalisation was performed on GAPDH housekeeping gene expression and then compared to control samples (DMSO-treated samples, Ctrl). The following primers were used: *GAPDH* Fw: 5’-CAACTACATGGTTTACATGTTC-3’, Rv: 5’-GCCAGTGGACTCCACGAC-3’; *ZNF598* Fw: 5’-GCTCATCCAGTCCATCAGGG-3’, Rv: 5’-GCAGGACCAGCAGCTCATTA-3’.

### Live-cell imaging analysis

To monitor red-Keima puncta, cells were plated (7*10^4^/well) onto a 12-well glass-bottom plate (MatTek) previously coated with 5μg/mL fibronectin, cultured for 24 hours in presence of 500nM Mps1i (or DMSO for Ctrl samples) or treated with Bafilomycin A_1_ as indicated in figure legend; for ZNF598 knock- down samples, see the protocol above. Cells were stained with 2μg/mL hoechst 33342 (Cell Signaling Technology) for 10 minutes and then imaged at 37° and 5% CO2 with a Leica TCS SP8 microscope with Acousto-Optical Beam Splitter (AOBS) equipped with a 405 diode, a 561 DPSS, a 640 HeNe, Argon laser lines and equipped with both Leica hybrid (HyD) and photomultiplier (PMT) detectors. Cells were imaged using a 63X/1.4NA oil objective. Green-Keima was excited with the 488nm Argon laser, red-Keima was excited with the 561nm laser line, and Hoechst with the 405nm diode. All emitting signals were detected with the HyD detectors. Acquisitions were performed using the Leica LasX software (version 3.1.5.16308). 12 independent image stacks were acquired in a 8-bit format every 2μm within a z-range of 10μm. The image pixel size was set to 90nm. Image analysis was performed with a custom pipeline using Fiji software(*60*). In particular, the average intensity of red-Keima puncta in a single image was calculated on the maximum intensity z-projection by dividing the raw integrated density inside manually segmented cells by the total cell area. Manual cell segmentation was performed by taking advantage of both Red- and Green-Keima signals. Red-Keima intensity was finally normalised by the average of the respective control sample and plotted with GraphPad PRISM (version 9.3.1).

To quantify RFP/GFP ratio with the fluorescent tandem construct RPS3-RFP-GFP (24-hour samples), cells were plated (5*10^4^/well) onto a 12-well glass-bottom plate (MatTek) previously coated with 5μg/mL fibronectin. 24 hours before imaging, 500nM Mps1i (or DMSO) were added and, then, cells were treated with Bafilomycin A_1_ as indicated in figure legend. For 72-hour samples, cells were pulsed for 24 hours with 500nM Mps1i (or DMSO for Ctrl) and, after drug washout, they were cultured for another 48 hours before imaging. After the corresponding treatments, all cells were stained with 2μg/mL hoechst 33342 (Cell Signaling Technology) for 10 minutes. Cells were imaged at 37° and 5% CO2 with a Leica TCS SP8 microscope with AOBS using a 63X/1.4NA oil objective. RFP was excited with the 561nm laser line, GFP with the 488 Argon laser and Hoechst with the 405nm diode. All emitting signals were detected with the HyD detectors. Acquisitions were performed using Leica LasX software. 30 independent image stacks were acquired in a 12-bit format every 0.5μm within a z-range of 2μm. The image pixel size was set to 180nm. Three biological replicas were performed. Image analysis was performed with a custom pipeline using Fiji software(*60*). Firstly, the ratio between RFP and GFP channels was calculated on a pixel basis on the maximum intensity z-projection of the image. Then, after a 2x2 average binning, the average ratio for all pixels within the same field of view was calculated. Pixels were not included in the average if (i) localized inside nuclei (whose masks were evaluated with a thresholding segmentation of the Hoechst channel), (ii) the ratio was less than 2 and (iii) the GFP value was 0 (i.e., not meaningful ratio). RFP/GFP ratios were finally normalised by the average of the respective control sample and plotted with GraphPad PRISM.

### Correlative light electron microscopy (CLEM) analysis

To visualise ribosomes in the lysosomal structures of aneuploid cells, correlative light electron microscopy (CLEM) technique was used. RPE1 Ribo-Keima cells were pulsed for 24 hours with 500nM Mps1i. After another 24-hours, cells were trypsinised and re-plated (1,5*10^4^/dish) onto 35mm gridded dishes (MatTek) previously coated with 5μg/mL fibronectin and cultured for another 24 hours (to reach the end 72-hour time-point). Before live-cell acquisitions, cells were stained with 2μg/mL hoechst 33342 (Cell Signaling Technology) for 10 minutes. Cells were then imaged with a Leica SP5 confocal microscope with incubator (37°C with 5% CO_2_) and LasAF software (version 3.1.5.16308), with 20X and 63X magnification objectives, 561nm (red-Keima channel) and 405nm laser excitation (hoechst channel) and 8μm Z size. Right after acquisition, cells were fixed adding the pre-heated fixative buffer (2,5% Glutaraldehyde in 0,1M cacodylate buffer, pH=7.4) for 1 hour at room temperature and washed in 0,1M cacodylate buffer. Samples were post-fixed in 1% OsO_4_, 1.5% K_4_Fe(CN)_6_, 0,1M sodium cacodylate for 1 hour at 4C° protected from light and washed. Then, samples were stained with Uranyl acetate 0,5% at 4C° overnight, protected from light. After washings with ddH_2_O, samples were dehydrated using ethanol 30% - 50% - 70% - 80% - 90% - 96%, and three steps in 100% ethanol. Samples were covered with a mixture of ethanol and epoxy resin 1:1 for 2 hours on a shaker at room temperature. Then, after two changes of pure epoxy resin, samples were embedded in fresh epoxy resin and polymerised overnight at 45°C plus 24 hours at 60°C. Embedded samples were then sectioned (70nm) on a Leica EM UC6 ultramicrotome, identifying back the cells of interest, previously acquired with fluorescence microscopy, using the reference coordinate system on the MatTek chamber. On a 300 Mesh grid, sections were contrasted with 2% aqueous uranyl acetate and stained with Sato’s lead stain for 2 minutes. After grid wash with pure water and drying, images were collected using a FEI Tecnai-12 transmission electron microscope. TEM and confocal images were aligned and merged to identify the objects highlighted by the red-Keima signal using Fiji software(*60*).

### Fluorescence-activated cell sorting (FACS) analysis

To measure the amount of misfolded and unfolded proteins in aneuploid cells, the cell-permeable fluorescent dye tetraphenylethene maleimide (TMI(*44*), kindly provided by Prof. Yuning Hong and Prof. Danny Hatters) was used. RPE1 cells were pulsed for 24 hours with 500nM Mps1i (or DMSO for Ctrl). After drug wash-out and 48 hours of culturing (or 24h hours after AZC treatment), cells were trypsinised and 1*10^6^ cells were resuspended and incubated in 50μM TMI in 1XPBS (45 minutes at 37°C). After being washed and resuspended in 1mL 1XPBS in FACS tubes, cells were analysed on a FACS Celesta 2 B-V-YG flow cytometer (BD Biosciences). 50.000 events per sample were collected with a 405nm laser and BV421 450/40nm bandpass filter. 2,5ug/mL propidium Iodide (Sigma-Aldrich) was used to select live cells. Data were analysed with FlowJo software (version 10.5.3). The same gate was applied to all samples to identify singlets and the threshold for TMI fluorescence was set on the negative control (not stained), to select per each sample cell containing higher-than-background TMI signal. The obtained TMI mean fluorescence intensities were plotted as Mps1i/Ctrl ratio, using GraphPad PRISM (version 9.3.1).

To analyse the distribution of RFP/GFP ratio in HCT116 RPS3-RFP-GFP cells, cells were plated (2,5*10^5^/well) onto a 6-well plate and, 24 hours before FACS analysis, they were pulsed with 500nM Mps1i (or DMSO for Ctrl) or treated with Bafilomycin A_1_, as indicated in figure legend. For 72-hour samples, cells were pulsed for 24 hours with 500nM Mps1i (or DMSO for Ctrl) and, after drug washout, they were cultured for another 48 hours before FACS analysis. Afterwards, samples were trypsinised, 1*10^6^ cells were washed in bovine serum albumin (BSA) 1% in 1XPBS and resuspended in 300uL 1XPBS in FACS tubes. FACS was carried out on a FACSMelody (BD Biosciences) and 30.000 events per sample were analysed with 561nm sand 488nm lasers. Data were processed with FlowJo software (version 10.5.3), by creating the new derived parameter RFP/GFP (561/488nm ratio) and normalising each RFP/GFP value to the mean value of the respective Ctrl sample. RFP/GFP distribution in each sample was visualised on histogram and data from replicates were represented on graphs as RFP/GFP change respect to the lower Ctrl value, using GraphPad PRISM (version 9.3.1).

### Autophagosome isolation and MS/MS analysis

Autophagosome isolation from aneuploid and pseudo-diploid cells was performed after about 3 cell cycles from drug treatments (500nM Mps1i or DMSO) and carried out as previously described(*35*). For this study, data derived from the most enriched functional annotations were considered as hints to proceed with further *in vitro* analysis.

### In vivo experiments with Drosophila melanogaster

Wing imaginal disc and eye-antennal imaginal disc were used to induce CIN and perform the screenings upon overexpression or depletion of Hsp70. Strains of *Drosophila melanogaster* were maintained on standard medium (4% glucose, 55 g/L yeast, 0.65% agar, 28 g/L wheat flour, 4 ml/L propionic acid and 1.1 g/L nipagin) at 25°C in light/dark cycles of 12 hours.

For CIN eye experiments, females carrying either the *ey-gal4* driver alone (5535, Bloomington Drosophila Stock Center) (control) or the *ey-gal4* driver together with the *UAS-bub3-RNAi* transgene (21037, Vienna Drosophila RNAi Center) (CIN) were crossed with males of the indicated genotypes (*UAS-GFP-RNAi, UAS-Hsp70A-RNAi, UAS-Hsp70B-RNAi or UAS-Hsp70A*) and progeny was kept at 25°C until they enclosed. The *ey-gal4* driver is expressed only in the eye primordium, and phenotypes were monitored in adult males and were represented on a graph as percentage of weak/strong, using GraphPad PRISM (version 9.3.1). To monitor cell death in wing imaginal disc, females carrying either the *ap-gal4*-driver expressing myristoylated-Tomato alone (*ap-gal4 UAS-myrT*, 32221, Bloomington Drosophila Stock Center) (control) or the *ap-gal4*-driver expressing myrT together with the UAS-bub3-RNAi transgene (21037, Vienna Drosophila RNAi Center) (CIN) were crossed with males of the indicated genotypes (*UAS-GFPi, UAS-Hsp70A-i, UAS-Hsp70B-i or UAS-Hsp70A*) and allowed to lay eggs on standard fly food for 24 hours at 25°C and kept at 25°C for another 24 hours. Then, larvae were switched to 29°C and maintained for 3 days before dissection. Experimental flies and control individuals were grown in parallel. The following transgenes were used for both experiments: *UAS-GFP-RNAi* (9331, Bloomington Drosophila Stock Center), *UAS-Hsp70A-RNAi* (42639, Bloomington Drosophila Stock Center), UAS-Hsp70B-RNAi (32997, Bloomington Drosophila Stock Center), *UAS-Hsp70A* (17624, Bloomington Drosophila Stock Center). To quantify cell death, immunostaining and imaging analysis were performed on wing imaginal discs of third instar larvae, by fixing them with formaldehyde 4% for 20 minutes and blocking for 1 hour in BBT (PBS + 0.1% Triton+ 0,3% BSA + 250mM NaCl). Primary antibodies were incubated overnight and with secondary antibodies for 2 hours. The following antibodies were used: anti-cleaved-Dcp1 (Cell Signaling Technology), Alexa 488-anti-rabbit secondary antibody (Jackson Immunoresearch). DAPI (ThermoFisher Scientific) was used to stain DNA, Zeiss LSM780 confocal microscope was used to image disc samples. The *ap-gal4* driver is expressed only in the dorsal (D) compartment of the wing primordium and size of the D compartment in the wing primordia was measured using Fiji software(*60*). Basal planes (delaminating-dying cells) were considered to determine the area positively labelled with Dcp-1 positive cells, and those values were normalised to the area of the dorsal region (D). Data were represented on a graph as cDcp1 signal / D area ratios, all of them normalised to the value obtained in the control (*UAS-GFP-RNAi*), using GraphPad PRISM (version 9.3.1).

### Polysome profiling analysis

Polysome profiles from aneuploid and control cells were obtained from RPE1 cells, plated to reach 70-80% confluency and pulsed 24 hours with 500nM Mps1i (or DMSO). After drug washout and 48-hour culturing (72-hour time-point), cells were treated with 10μg/mL cycloheximide for 3,5 minutes at 37°C. After three washes with cold PBS, 10μg/mL cycloheximide, cells were directly lysed in the dish with 300μL of ice-cold lysis buffer (10mM NaCl, 10mM MgCl_2_, 10mM Tris-HCl pH 7.5, 1% Triton-X100, 1% NaDeoxycholate, 0.2U/μL Rnase inhibitor, 1mM dithiothreitol, 10μg/ml cycloheximide, 0.005U/ μL DNAse I) and centrifuged for 5 minutes at 18000 g at 4°C. Cell lysates were carefully added over 10%–40% sucrose gradient prepared in : 30mM Tris-HCl pH 7.5, 100 mM NaCl, 10 mM MgCl_2_ in DEPC-treated water. Samples were ultracentrifuged in a SW41Ti rotor (Beckman) for 1 hour and 40 minutes at 180,000g at 4°C in a Beckman Optima LE-80K Ultracentrifuge. After stabilisation (20 minutes on ice), gradients were fractionated in 1mL volume fractions with continuous monitoring absorbance at 254nm using a Teledyne ISCO UA-6 UV/VIS detector. Proteins were isolated from the fractions using the methanol/chloroform method, as described in Lauria *et al.* (*61*) and used for Western blot analysis. Western blots were performed using the following primary antibodies: anti-RPL26 (Abcam), anti-RPS6 (Cell Signaling Technology), and the appropriate HRP-conjugated secondary antibodies (Santa Cruz Biotechnology). Detection was performed using the ECL Prime Western Blotting Detection Reagent (Amersham Biosciences). The fraction of ribosomes in polysomes (FRP) was calculated from polysomal profiles as the ratio between the area under the curve of polysomes and the area under the curve of polysomes plus the area of the 80S peak, as previously described(*62*). Polysome profiles from four different biological replicates were considered for each sample and the profiles of each biological replicate were obtained from three technical replicates.

### Data analysis of cancer cells

To analyse the link between heat shock response and aneuploidy, we used the CCLE data from the Dependency Map (DepMap) platform version 22q2 (www.DepMap.org). We compared the “highly aneuploid” cancer cell lines, defined as the top-quartile of the number of arm-level events (either chromosome gains or losses) (aneuploidy score ≥ 21), and “low aneuploid” cancer cell lines, as the bottom-quartile of the number of arm-level events (aneuploidy score ≤ 8). Aneuploidy scores were obtained from Zerbib *et al*(*54*). We analyzed both mRNA expression, RNAi dependency and CRISPR screens data for several heat shock genes (HSF1, HSP90B1, HSP90AB1, HSP90AA1, HSPA1B). Due to lack of RNAi data for the gene HSPA1B in the DepMap version used for this paper, we analyzed this specific gene in a newer version (23q4). For statistical analysis we performed a two-tailed Student’s t-test and plotted the results on a violin plot, using GraphPad PRISM (version 9.3.1).

To correlate ZNF598 expression in patient primary cancer samples, data from TCGA were analysed. The correlation between ZNF598 mRNA expression (in RSEM) and the aneuploidy score of the tumours was assessed by linear regression (Y=0,02X+9,66; Spearman ρ=0.2524), using GraphPad PRISM. Gene expression data were obtained from the CCLE using the DepMap 22q1 release. Single sample gene set enrichment analysis (ssGSEA) was performed using GenePatterns(*63*, *64*) and calculated for the following signatures: “KEGG_RIBOSOME”, “GOBP_RIBOSOME_ASSEMBLY”. Aneuploidy scores were obtained from Cohen-Sharir *et al.*(*65*) and cancer cells were divided into “highly aneuploid” and “low aneuploid”, based on their aneuploidy scores. Significance was calculated by an unpaired two-tailed Student’s t-test and the results were plotted on a violin plot, using GraphPad PRISM. To associate gene expression pathways with ZNF598 expression, pathway enrichment analysis was performed using PreRanked GSEA, and an enrichment analysis was performed on the top 200 negatively correlated genes with ZNF598 expression using the MsigDB database (https://www.gsea-msigdb.org/gsea/msigdb/). Shown are the results for the most negatively correlated genes looking at the signatures from the KEGG and Reactome datasets.

### Quantification and statistical analysis

To test statistical significance, experiments were performed in at least three biological replicates (otherwise indicated in figure legends). The statistical analysis was performed using GraphPad PRISM (version 9.3.1). Details of the statistical tests were reported in figure legends. Error bars are shown if n>2 and represent SEMs or SDs. When not indicated on the graphs, differences were not statistically significant. *P*-value is shown in the graphs where it applies (*p*<0.05 was considered significant), as follow: * for *p*<0,05, ** for *p*<0,01, *** for *p*<0,001 or **** for *p*<0,0001.

## Acknowledgments

The authors would like to thank members of the Santaguida lab for helpful and insightful discussions; Zuzana Storchová and Wade Harper for providing reagents. This work was supported by the Italian Association for Cancer Research (AIRC-MFAG 2018 - ID. 21665, AIRC-Bridge Grant 2023 - ID. 29228 and AIRCG-IG 2024 – ID. 31023 to S.S.), Ricerca Finalizzata (GR-2018-12367077 to S.S.), Fondazione Cariplo (S.S.), the Rita-Levi Montalcini program from MIUR (to S.S.) and the Italian Ministry of Health with Ricerca Corrente and 5x1000 funds (S.S.). M.R.I. was supported by Fondazione IEO-MONZINO ETS and by the Italian Association for Cancer Research (ID 26738-2021 and ID 31556-2024). Work in M.M.’s lab is funded by the BFU2016-77587-P grant from Ministerio de Ciencia e Innovación (Government of Spain). The Ben-David lab is supported by the European Research Council Starting Grant (grant #945674 to U.B.-D.), the Israel Cancer Research Fund (U.B.-D.), the Israel Science Foundation (grant #1805/21 to U.B.-D.), and the BSF project grant (grant #2019228 to U.B.-D.). A.P. is supported by the European Union - NextGenerationEU: National Center for Gene Therapy and Drug based on RNA Technology, CN3 - Spoke 3 (code: CN00000041; PNRR MUR – M4C2 – Action 1.4- Call “Potenziamento strutture di ricerca e di campioni nazionali di R&S”, CUP: B83C22002870006).

## Author contributions

Investigation, S.V., S.S., M.R.I, L.B., A.L., M.R.M., D.D., Y.E., L.P.V., J.A.P., M.M., A.P.; Funding acquisition and supervision: U.B.D., G.V., M.M. and S.S.; Conceptualization: S.V. and S.S.; Writing: S.V. and S.S. with input from all authors. All authors discussed the results and commented on the manuscript.

## Data and materials availability

All data are available in the main text or the supplementary materials.

