## Supplemental materials for "Ribosome Quality Control Mitigates Proteotoxic Stress in Aneuploid Cells"

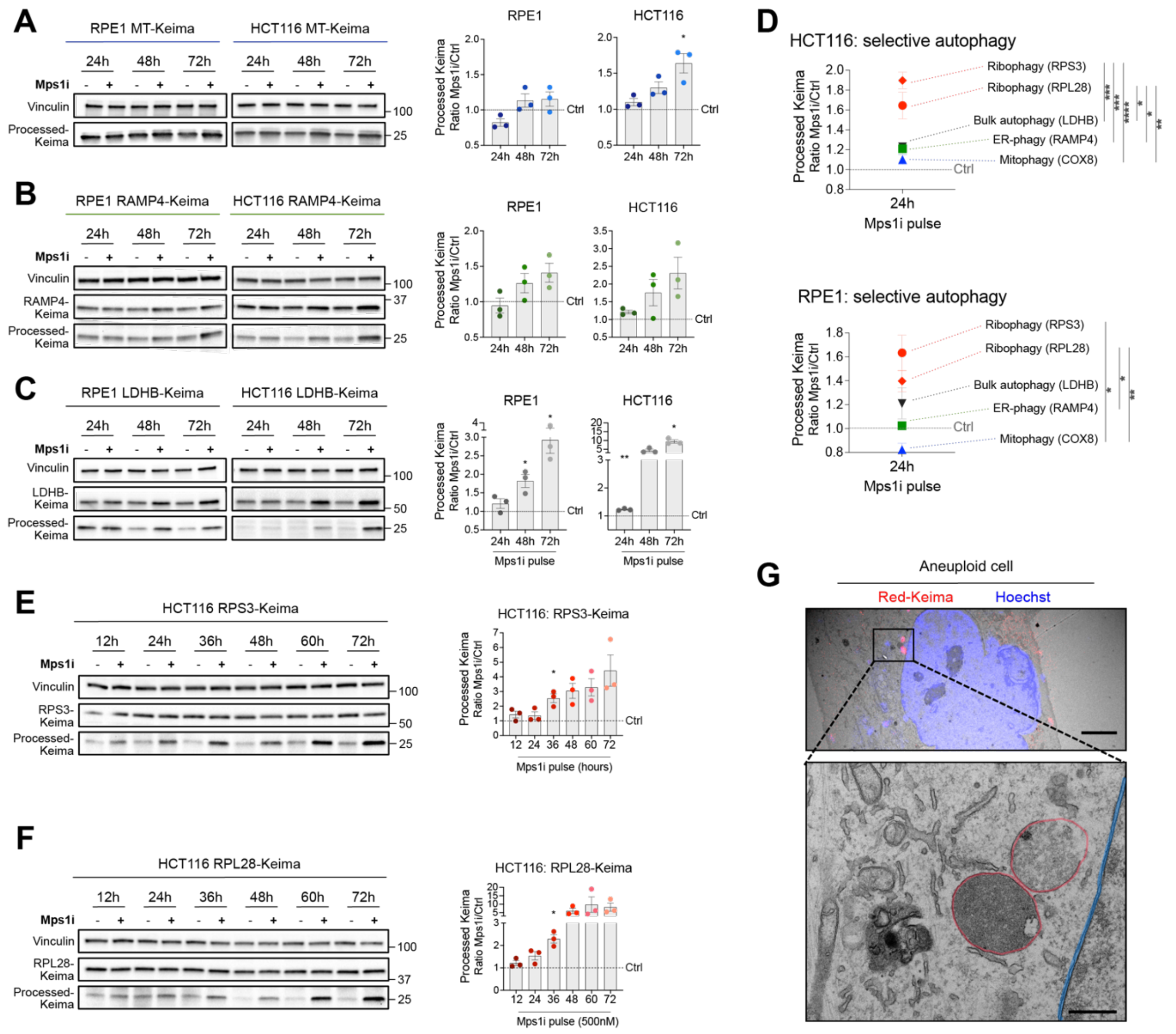


#### Figure S1

**Figure S1. Aneuploidy affects the autophagic removal of ribosomes.**

- **A.** Representative immunoblots and replicate quantitation of the indicated MT-Keima cell lines treated with Mps1i pulse or DMSO (control) and collected at 24, 48 or 72 hours; processed-Keima level is proportional to mitochondria degradation; vinculin was used as loading control. Mean ± SEM, $n$=3; one sample and Wilcoxon test (Mps1i at each time-point *vs* respective control=1): * indicates $p$=0.0419.
- **B.** Representative immunoblots and replicate quantitation of the indicated RAMP4-Keima cell lines treated with Mps1i pulse or DMSO (control) and collected at 24, 48 or 72 hours; increased level of processed-Keima indicates ER degradation; vinculin was used as loading control. Mean ± SEM, $n$=3; one sample and Wilcoxon test (Mps1i at each time-point *vs* respective control=1): *ns*.

**C.** Representative immunoblots and replicate quantitation of the indicated LDHB-Keima cell lines treated with Mps1i pulse or DMSO (control) and collected at 24, 48 or 72 hours; increased level of processed-Keima indicates increased bulk autophagy; vinculin used as loading control. Mean ± SEM, *n*=3; one sample and Wilcoxon test (Mps1i at each time-point *vs* respective control=1): * indicates *p*=0.0432 (RPE1-48h) or *p*=0.0332 (RPE1-72h) or *p*=0.0124 (HCT116-72h), ** indicates *p*=0.0081.

**D.** Plot of the processed-Keima levels of the indicated samples, to highlight the different behaviour of the analysed selective autophagies and bulk autophagy right after aneuploidy induction (24 hours of Mps1i/DMSO treatment). The graphed values are the same as those shown in Figure 1D-E, Figure S1A-C. Mean ± SEM at 24 hours; one-way ANOVA, followed by Tukey's multiple comparison test: * indicates *p*=0.0183 (HCT116 RPL28 *vs* LDHB) or *p*=0.0139 (HCT116 RPL28 *vs* RAMP4) or *p*=0.0162 (RPE1 RPS3 *vs* COX8) or *p*=0.0110 (RPE1 RPL28 *vs* RAMP4), ** indicates *p*=0.0033 (HCT116 RPL28 *vs* COX8) or *p*=0.0015 (RPE1 RPL28 *vs* COX8), *** indicates *p*=0.0003 (HCT116 RPS3 *vs* LDHB) or *p*=0.0003 (HCT116 RPS3 *vs* RAMP4), **** indicates *p*<0.0001.

**E.** Representative immunoblot and replicate quantitation of processed-Keima levels in HCT116 RPS3-Keima cells treated with Mps1i pulse or DMSO (control) and collected at 12, 24, 36, 48, 60 or 72 hours; vinculin was used as loading control. Mean ± SEM, *n*=3; one sample and Wilcoxon test (Mps1i at each time-point *vs* respective control=1): * indicates *p*=0.0360.

**F.** Representative immunoblot and replicate quantitation of processed-Keima levels in HCT116 RPL28-Keima cells treated with Mps1i pulse or DMSO (control) and collected at 12, 24, 36, 48, 60 or 72 hours; vinculin was used as loading control. Mean ± SEM, *n*=3; one sample and Wilcoxon test (Mps1i at each time-point *vs* respective control=1): * indicates *p*=0.0211.

**G.** Representative images of RPE1 Ribo-Keima cell treated with Mps1i pulse and selected for CLEM: alignment and merge between confocal image and electron microscopy image (top) and magnified electron microscopy images showing ribosomes within Red-Keima positive single-membraned autolysosomes (red profiles, bottom); hoechst has been used to stain DNA (blue profiles); scale bar 5µm (top) and 500nm (bottom).

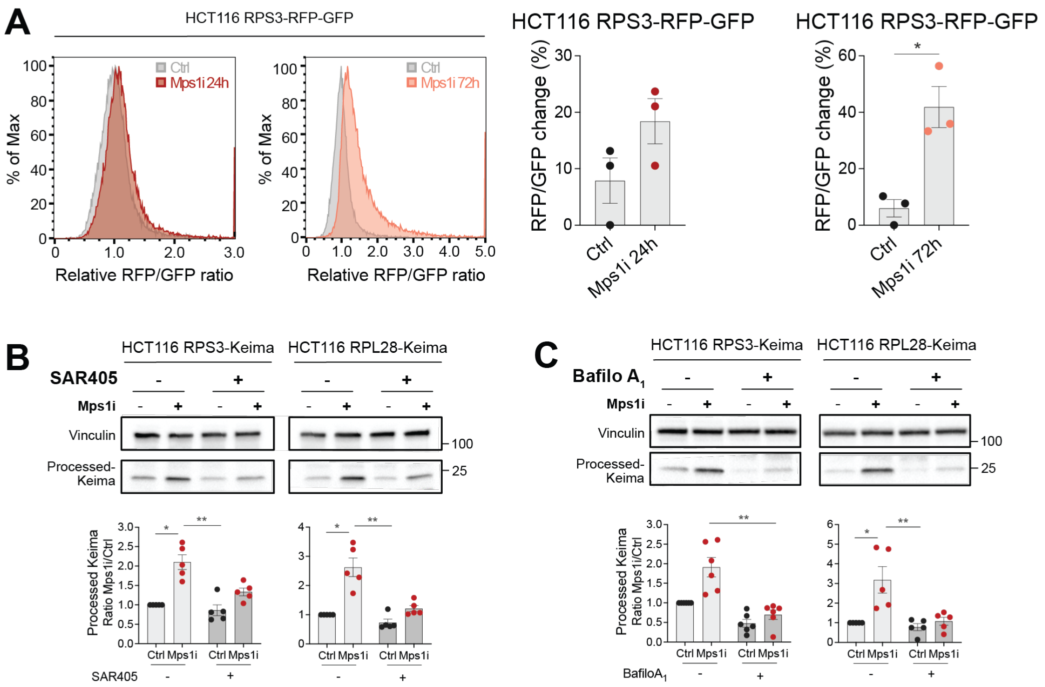


### Figure S2

**Figure S2. Ribosome clearance in aneuploid cells takes place through canonical lysosome-mediated degradation.**

**A.** Representative FACS analysis and replicate quantitation of HCT116 RPS3-RFP-GFP cells, treated with Mps1i pulse or DMSO (Ctrl) and collected at 24 or 72 hours. Relative frequency distributions of RFP/GFP ratio are shown and replicates have been plotted as % change of RFP/GFP ratio. Mean± SEM, *n*=3; unpaired Student's t-test: * indicates *p*=0.0106.

**B.** Representative immunoblots and replicate quantitation of processed-Keima levels in the indicated HCT116 Ribo-Keima cell lines treated with Mps1i pulse or DMSO (control), collected at 24 hours and upon SAR405 treatment (1μM, 24h); vinculin was used as loading control. Mean ± SEM, *n*=5; Kruskal-Wallis test, followed by Dunn's multiple comparison test: * indicates *p*=0.0218 (HCT116 RPS3-Keima) or *p*=0.0183 (HCT116 RPL28-Keima), ** indicates *p*=0.0023 (HCT116 RPS3-Keima) or *p*=0.0018 (HCT116 RPL28-Keima).

**C.** Representative immunoblots and replicate quantitation of processed-Keima levels in the indicated HCT116 Ribo-Keima cell lines treated with Mps1i pulse or DMSO (control), collected at 24 hours and upon Bafilomycin $A_1$ (Bafilo$A_1$) treatment (100nM, 6h); vinculin was used as loading control. Mean ± SEM, *n*=6 for HCT116 RPS3-Keima, *n*=5 for HCT116 RPL28-Keima; Kruskal-Wallis test, followed by Dunn's multiple comparison test: * indicates *p*=0.0242 (HCT116 RPL28-Keima), ** indicates *p*=0.0052 (HCT116 RPS3-Keima) or *p*=0.0089 (HCT116 RPL28-Keima).

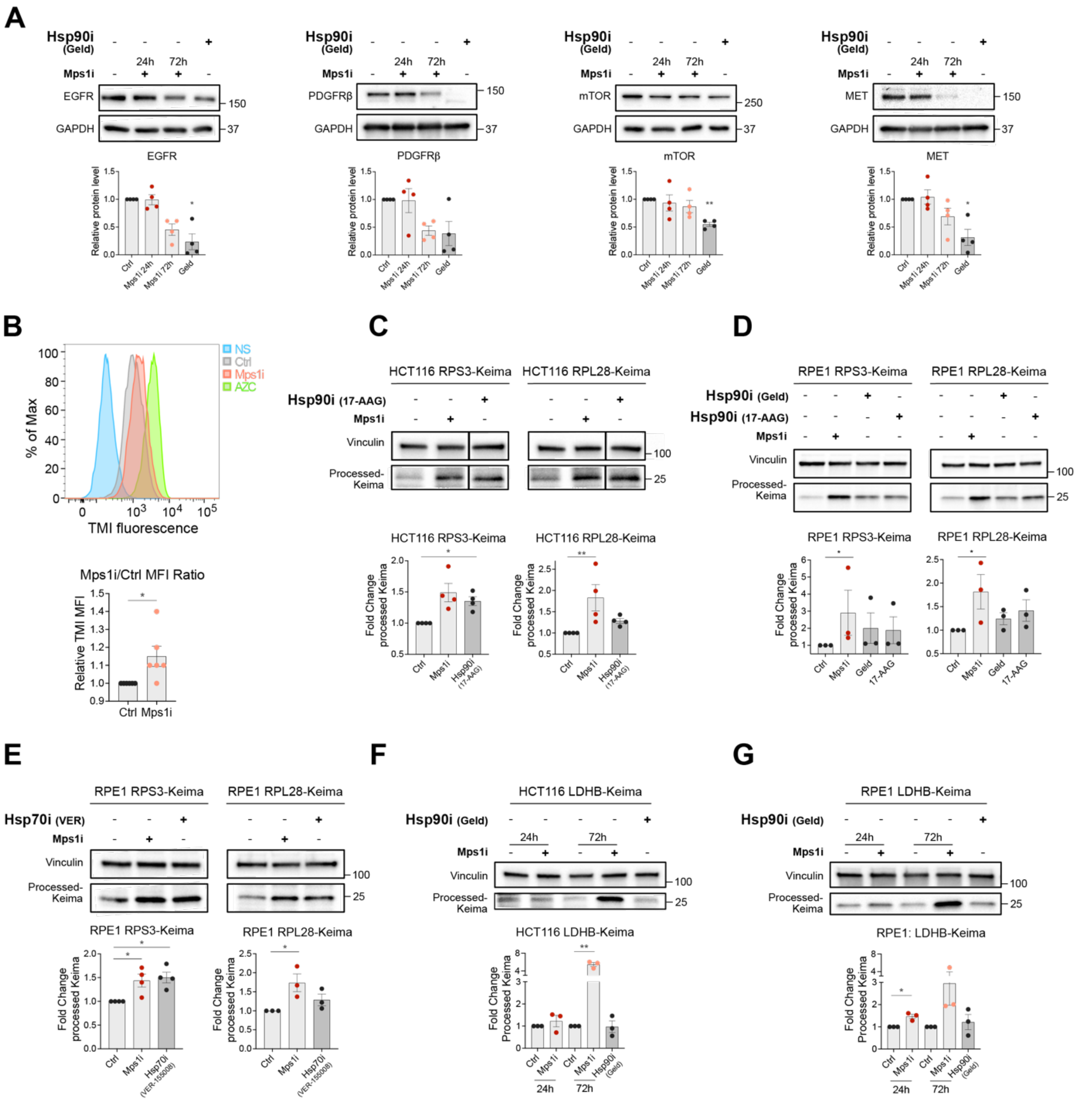


### Figure S3

**Figure S3. Protein folding impairment affects the autophagic removal of ribosomes.**

**A.** Representative immunoblots and replicate quantitation of the indicated Hsp90 chaperone clients in RPE1 cells treated with Mps1i pulse or DMSO (control) and collected at 24 or 72 hours; Hsp90i Geldanamycin (1µM, 12h) was used as a positive control; GAPDH was used as loading control. Mean ± SEM, *n*=4; Kruskal-Wallis test, followed by Dunn's multiple comparison test: * indicates *p*=0.0263 (EGFR) or *p*=0.0134 (MET), ** indicates *p*=0.0082 (mTOR).

**B.** Representative FACS analysis and replicate quantitation of tetraphenylethene maleimide (TMI) fluorescence in RPE1 cells treated with Mps1i pulse or DMSO (Ctrl) and collected at 72 hours; not-stained cells (NS) were used as negative control and L-azetidine-2-carboxylic acid (AZC)-treated cells (10mM, 24h) were used as positive control. Mean ± SEM, *n*=6; one sample and Wilcoxon test: * indicates *p*=0.0446.

**C.** Representative immunoblots and replicate quantitation of processed-Keima levels in the indicated HCT116 Ribo-Keima cell lines treated for 24 hours with Mps1i, DMSO (control) or Hsp90i 17-AAG (1µM); vinculin has been used as loading control. Mean ± SEM, *n*=4; Kruskal-Wallis test, followed by Dunn's multiple comparison test: * indicates *p*=0.0189, ** indicates *p*=0.0064.

**D.** Representative immunoblots and replicate quantitation of processed-Keima levels in the indicated RPE1 Ribo-Keima cell lines treated for 24 hours with Mps1i, DMSO (control), Hsp90i Geldanamycin (1µM) or Hsp90i 17-AAG (1µM); vinculin has been used as loading control. Mean ± SEM, *n*=3; Kruskal-Wallis test, followed by Dunn's multiple comparison test: * indicates *p*=0.0258 (RPE1 RPS3-Keima) or *p*=0.0396 (RPE1 RPL28-Keima).

**E.** Representative immunoblots and replicate quantitation of processed-Keima levels in the indicated RPE1 Ribo-Keima cell lines treated for 24 hours with Mps1i, DMSO (control) or Hsp70i VER-155008 (50µM); vinculin was used as loading control. Mean ± SEM, *n*=4 for RPE1 RPS3-Keima and *n*=3 for RPE1 RPL28-Keima; Kruskal-Wallis test, followed by Dunn's multiple comparison test: * indicates *p*=0.0434 (RPE1 RPS3-Keima Ctrl *vs* Mps1i) or *p*=0.0252 (RPE1 RPS3-Keima Ctrl *vs* Hsp70i) or *p*=0.0199 (RPE1 RPL28-Keima Ctrl *vs* Mps1i).

**F.** Representative immunoblots and replicate quantitation of processed-Keima levels in HCT116 LDHB-Keima cells treated with Mps1i pulse or DMSO (control) and collected at 24 or 72 hours, or Hsp90i Geldanamycin (1µM); vinculin was used as loading control. Mean ± SEM, *n*=3; one sample and Wilcoxon test (Mps1i 24h and Geld *vs* respective control=1): *ns*; one sample and Wilcoxon test (Mps1i 72h *vs* respective control=1): ** indicates *p*=0.0021.

**G.** Representative immunoblots and replicate quantitation of processed-Keima levels in RPE1 LDHB-Keima cells treated with Mps1i pulse or DMSO (control) and collected at 24 or 72 hours, or Hsp90i Geldanamycin (1µM); vinculin was used as loading control. Mean ± SEM, *n*=3; one sample and Wilcoxon test (Mps1i 24h and Geld *vs* respective control=1): * indicates *p*=0.0370; one sample and Wilcoxon test (Mps1i 72h *vs* respective control=1): *ns*.

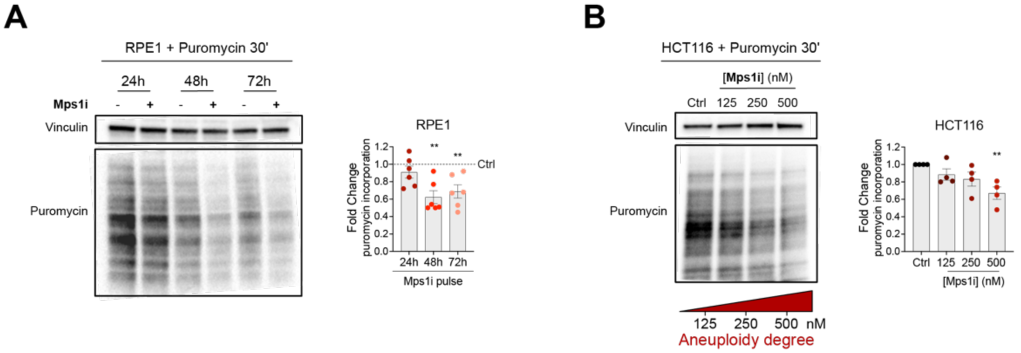


#### Figure S4

**Figure S4. Increasing aneuploidy-associated stresses and aneuploidy degree correlate with increasing autophagic removal of ribosomes.**

**A.** Representative immunoblot and replicate quantitation of puromycin incorporation (10μg/mL over 30 minutes) into RPE1 cells treated with Mps1i pulse or DMSO (control) and collected at 24, 48 or 72 hours; vinculin was used as loading control for puromycin smear quantitation. Mean ± SEM, $n$=6; one sample and Wilcoxon test (Mps1i *vs* respective control=1): ** indicates $p$=0.0036 (48h) or $p$=0.0093 (72h).

**B.** Representative immunoblot and replicate quantitation of puromycin incorporation (10μg/mL over 30 minutes) into HCT116 cells treated for 24 hours with 125nM, 250nM or 500nM Mps1i or DMSO (Ctrl); vinculin was used as loading control for puromycin smear quantitation. Mean ± SEM, $n$=4; Kruskal-Wallis test, followed by Dunn's multiple comparison test: ** indicates $p$=0.0093.

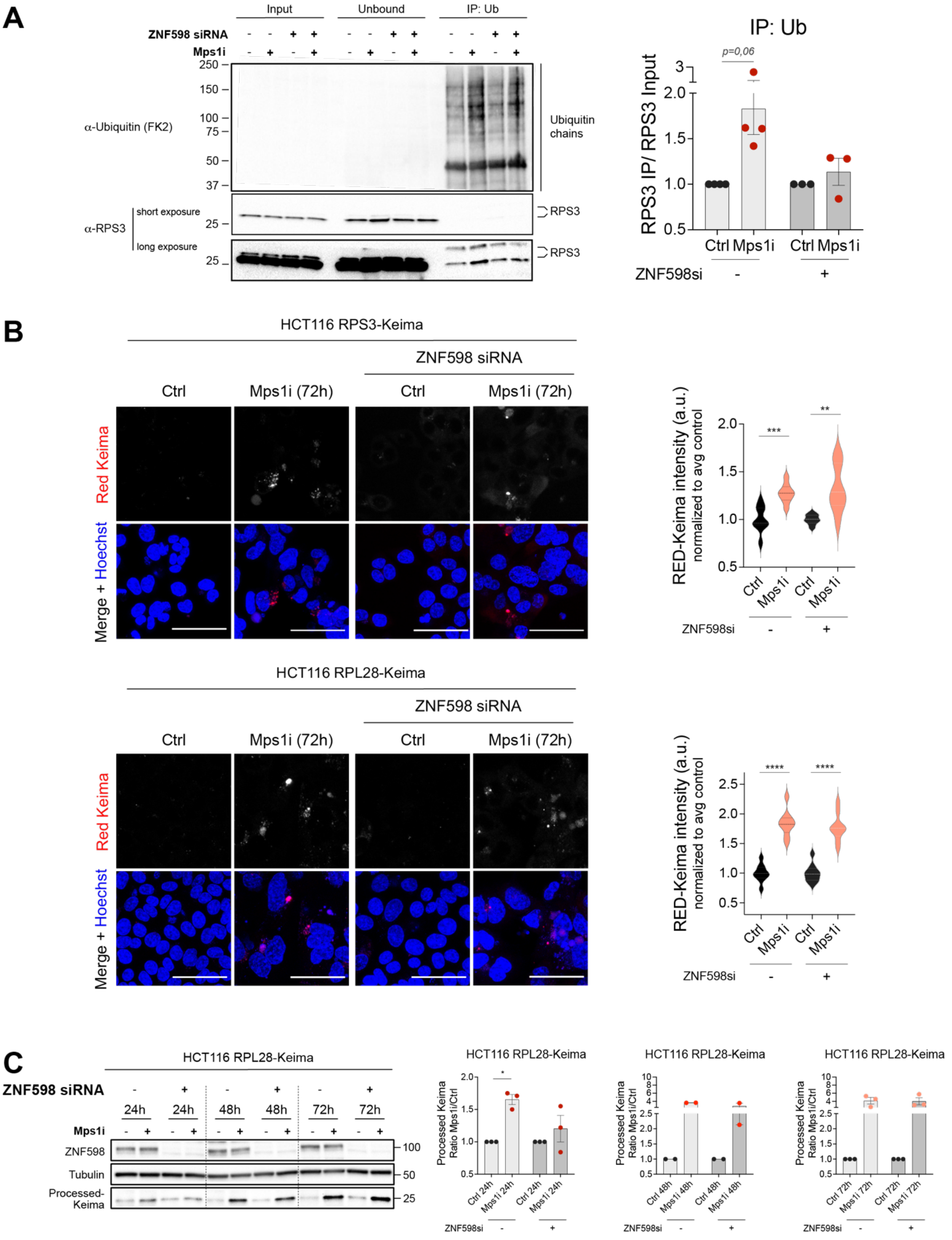
A
Input
Unbound
IP: Ub
ZNF598 siRNA
Mps1i
α-Ubiquitin (FK2)
Ubiquitin chains
α-RPS3
short exposure
long exposure
RPS3
250
150
100
75
50
37
25
IP: Ub
p=0,06
RPS3 IP/ RPS3 Input
Ctrl
Mps1i
ZNF598si
B
HCT116 RPS3-Keima
ZNF598 siRNA
Ctrl
Mps1i (72h)
Red Keima
Merge + Hoechst
RED-Keima intensity (a.u.) normalized to avg control
HCT116 RPL28-Keima
C
HCT116 RPL28-Keima
ZNF598 siRNA
24h
48h
72h
Mps1i
ZNF598
Tubulin
Processed-Keima
Processed Keima Ratio Mps1i/Ctrl
ZNF598si

**Figure S5**

**Figure S5. The E3-ligase ZNF598 ubiquitylates ribosomes and mediates their autophagic removal in aneuploid cells.**

**A.** Representative immunoblot and replicate quantitation of ubiquitin (Ub) immunoprecipitation (IP) experiment on HCT116 cells upon ZNF598 siRNA (or non-targeting siRNA) and treated for 24 hours with Mps1i or DMSO (control); RPS3 was blotted to assess its level of ubiquitylation, short and long exposures of the same membrane are shown. Mean ± SEM, *n*=4 for non-targeting siRNA samples, *n*=3 for ZNF598 siRNA samples; one sample and Wilcoxon test (Mps1i *vs* respective control=1): *p*=0.06; unpaired Student's t-test (Mps1i *vs* ZNF598siRNA Mps1i): *ns*.

**B.** Representative live-cell images and replicate quantitation of indicated HCT116 Ribo-Keima cells lines upon ZNF598 siRNA (or non-targeting siRNA), treated with Mps1i pulse or DMSO (Ctrl) and analysed at 72 hours. Red-Keima intensity is proportional to ribosome degradation; Hoechst has been used to stain DNA; scale bars, 50µm. Upper quartile, lower quartile and median of each violin plot are shown, *n*=9 fields of view; unpaired Student's t-test: ** indicates *p*=0.0013, *** indicates *p*=0.0004, **** indicates *p*<0.0001.

**C.** Representative immunoblots and processed-Keima quantitation of HCT116 RPL28-Keima cells upon ZNF598 siRNA (or non-targeting siRNA), treated with Mps1i pulse or DMSO (control) and collected at 24, 48 or 72 hours; ZNF598 was blotted as knock-down control and tubulin used as loading control. Mean ± SEM, *n*=3 for 24- and 72-hour samples, *n*=2 for 48-hour samples; one sample and Wilcoxon test (each Mps1i *vs* respective control=1): * indicates *p*=0.0138, unpaired Student's t-test (each Mps1i *vs* Mps1i+ZNF598siRNA): *ns*.

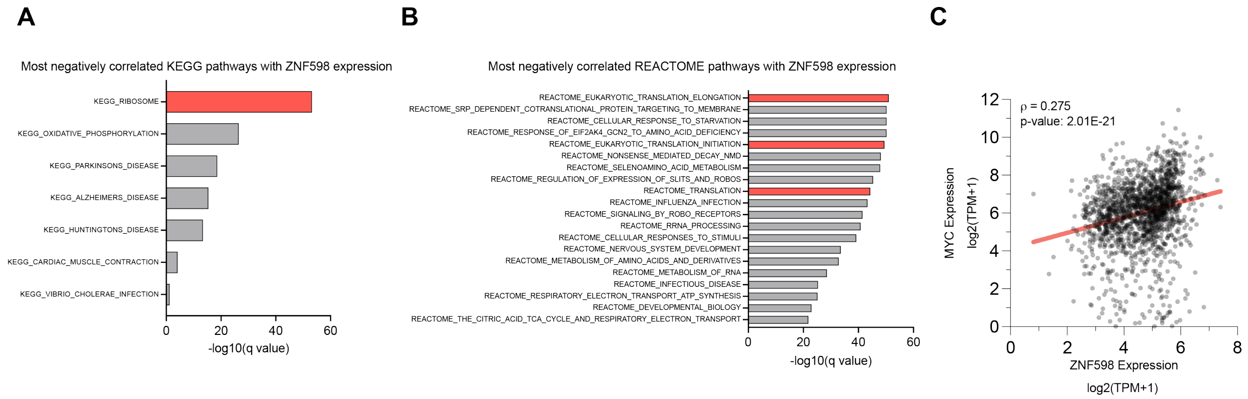


**Figure S6**

**Figure S6. Ribosomal and translational signatures anticorrelate with ZNF598 expression.**

**A.** Top 7 KEGG terms among the most negatively correlated with ZNF598 mRNA expression levels. X axis presents statistical significance (–$\log_{10}$ (q-value)). The red bar highlights the signature of interest.

**B.** Top 20 REACTOME terms among the most negatively correlated with ZNF598 mRNA expression levels. X axis presents statistical significance (–$\log_{10}$ (q-value)). Red bars highlight the signature of interest.

**C.** Correlations between the mRNA expression levels of ZNF598 and c-Myc in cancer cell lines. The trend line for Spearman's correlation is in red: ρ=0,275, *p*=2.01E-21.
